# Peroxisome protein import deficiency causes heart failure in mouse and human

**DOI:** 10.1101/2025.10.13.681307

**Authors:** Julia Hofhuis, Malte Tiburcy, Yelena Hartmann, Kristina Bersch, Ahmed Wagdi, Óscar Gutiérrez-Gutiérrez, Richard Solano, Leonie Thiele, Maya Walper, Mona Göbel, Wolfgang Hübner, Branimir Berecic, Marilù Casini, Petros Tirilomis, Andreas Unger, Noa Lipstein, Samuel Sossalla, Thomas Huser, Katrin Streckfuss-Bömeke, Manar Elkenani, Karl Toischer, Michael Kohlhaas, Jan Dudek, Wolfram-Hubertus Zimmermann, Niels Voigt, Tobias Bruegmann, Christoph Maack, Lukas Cyganek, Sven Thoms

## Abstract

Peroxisomes are ubiquitous cellular organelles with potentially vital roles in lipid and reactive oxygen metabolism. The metabolic demands of the heart are substantial; however, the contribution of peroxisomes to cardiac development, health, and their role in heart failure (HF) remain largely unexplored. We developed and examined a mouse and an engineered human myocardium (EHM) model with a deficiency in cardiac peroxisome biogenesis to investigate the role of peroxisomes in cardiac function and pathology. In the EHM, loss of peroxisome protein import and subsequent peroxisomal metabolic impairment trigger mitochondrial damage and compromise cellular respiration and energy production. Peroxisome dysfunction results in incoherent electrical conduction, defective Ca^2+^-handling, and ultimately presentation of a HF phenotype with pathological force generation. These phenotypes are mirrored in an orthogonal murine model system with defective cardiac peroxisome biogenesis. Preload-dependent deficits in force generation due to insufficient energy supply are eventually fatal. Thus, peroxisomes play an important role in sustaining normal heart operations. Vice versa, peroxisome maintenance is compromised in pressure overload-induced HF, establishing peroxisomes as potential modulators of pathology and targets of therapy.

## Introduction

Peroxisomes are essential organelles present in all nucleated cells, distinguished by their hydrogen peroxide-generating oxidases and hydrogen peroxide-degrading catalases^1^. Their significance in human cellular homeostasis stems from their involvement in a wide array of critical metabolic functions^2^. These include lipid metabolism, the management of reactive oxygen species (ROS), and the processing of various intermediate products of secondary metabolism. Working alongside mitochondria, peroxisomes degrade fatty acids through β-oxidation and serve as the exclusive site for α-oxidation of branched-chain fatty acids. Further, peroxisomes are essential for the synthesis of key biomolecules such as plasmalogens.

While peroxisomes have been extensively studied in the liver, kidneys, and brain, emerging research highlights their roles beyond metabolism^2,3^. Defects in peroxisome formation, classified as peroxisome biogenesis disorders (PBDs), have profound consequences for cellular function and human health^4^. These disorders arise from mutations in genes responsible for importing matrix and membrane proteins into the peroxisome. Among these, PEX5 plays a crucial role as a soluble receptor for peroxisomal targeting signal type 1 (PTS1)-containing proteins, mediating their recognition, docking, and translocation into peroxisomes^3,5^. A deficiency in PEX5 leads to the destabilization of peroxisomal proteins in the cytosol, compromising nearly all peroxisomal functions. Clinically, PBDs present as severe multi-organ diseases, with predominant neuro-sensory and developmental manifestations. Zellweger syndrome, the prototypical and most severe form of PBD, is often fatal within the first year of life. In rarer cases however, PBD can manifest as a life-threatening condition leading to heart failure (HF)^6^, underscoring the critical role of peroxisomes in maintaining cellular and systemic health.

HF is a severe and progressive condition characterized by the heart’s inability to pump blood efficiently. At the molecular level, its pathogenesis is driven by disruptions in excitation-contraction (EC) coupling, impaired contractility, and dysregulated Ca^2+^ handling. A central factor in this progressive decline is the inability of cardiac mitochondria to meet the high energy demands required for sustained muscle contraction and relaxation^7^. In fact, in various etiologies of HF, such mechano-energetic uncoupling, mediated either by primary defects in EC coupling or mitochondrial energetics, plays an important role in the development and progression of HF^8^. Despite the ubiquitous presence of peroxisomes and their well-established role in energy metabolism, their functions in both healthy and diseased cardiac tissue are largely unknown.

## Results

### A human induced pluripotent stem cell-derived cardiomyocyte model of peroxisome protein import deficiency

To investigate human cardiac peroxisomes, we developed pluripotent stem cell and tissue models. We decided to target PEX5, a pivotal component in the peroxisomal protein import pathway. Interfering with PEX5 function affects the import of all peroxisomal matrix proteins, retaining them in the cytosol, where they are disconnected from their metabolic pathways and become targets of protein degradation. In a previous study, we identified and characterized the homozygous stop codon mutation Q133X in a patient with severe Zellweger phenotype who died in early childhood^9^. Using CRISPR/Cas9 and homology-directed repair, we generated human *PEX5*-deficient iPSCs (*PEX5^iKO^*) by insertion of Q133X leading to a truncation of *PEX5* in exon 5 (Figure 1A-C). Gene editing was confirmed by Sanger Sequencing (Figure 1D), and *PEX5*-deficient hiPSCs lines were characterized for pluripotency and genomic stability. *PEX5^iKO^* displayed human stem cell morphology (Figure 1E), and showed robust expression of key pluripotency markers OCT4 (Figure 1F,H), TRA1-60 (Figure 1G,I), and NANOG (Figure 1J). Molecular karyotyping using a genome-wide SNP array demonstrated chromosomal stability of hiPSCs after genome editing (Figure 1K).

**Figure 1:**
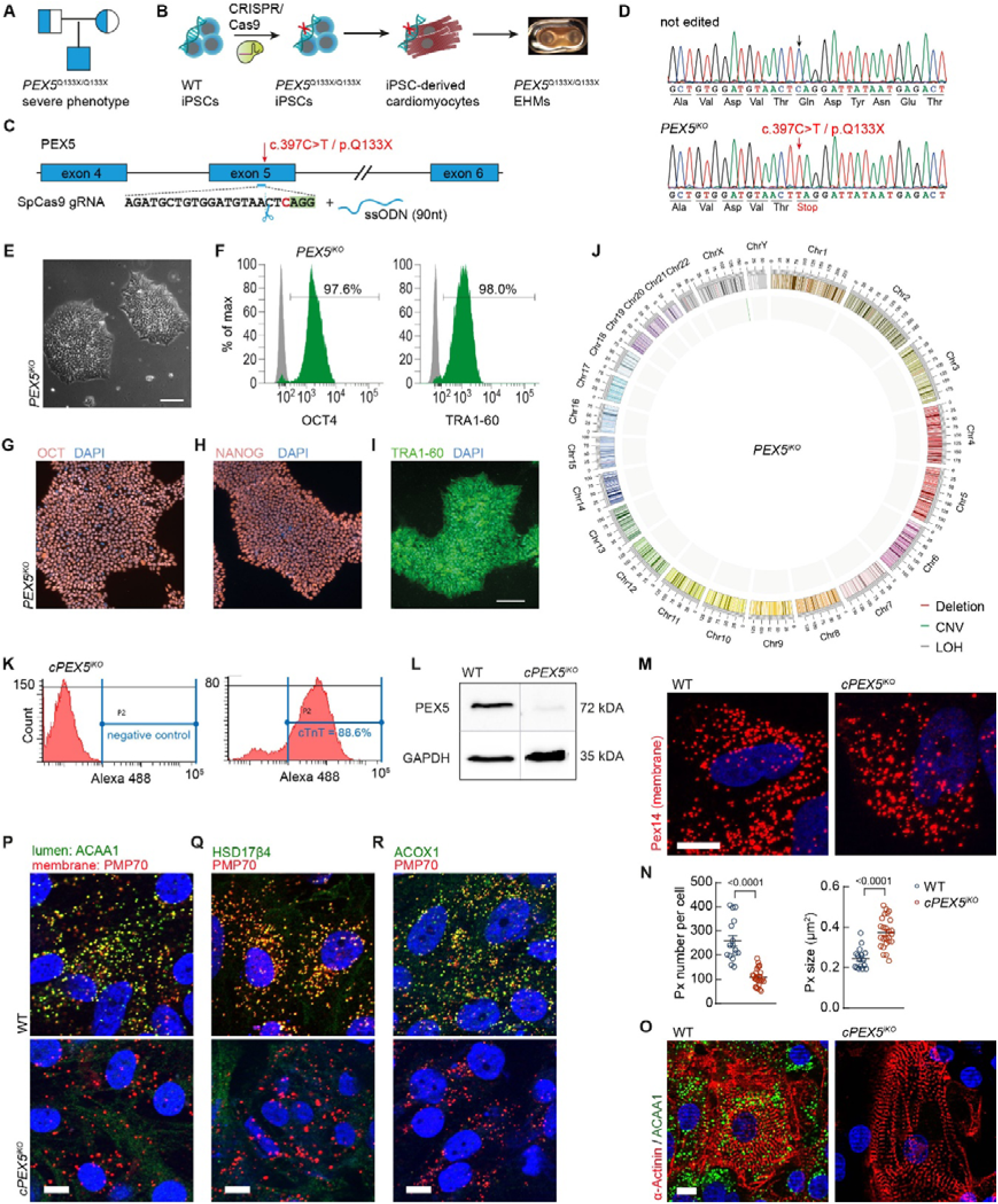
Modelling peroxisome deficiency in human iPSC cardiomyocytes. **(A)** The homozygous point mutation Q133X leads to a severe phenotype with death in the first years of life. (**B,C**) Generation of *PEX5*-deficient hiPSCs (*PEX5^iKO^*) by CRISPR/Cas9-based insertion of the pathological variant Q133X. *PEX5^iKO^*were differentiated into hiPSC-CMs (*cPEX5^iKO^*) and EHM. (**D**) Successful insertion of the variant in homozygous state was verified by sequencing. (**E**) *PEX5^iKO^* showed typical human stem cell-like morphology; scale: 100 µm. (**F**) Pluripotency analysis of OCT4 and TRA-1-60 assessed by flow cytometry; grey peaks represent negative control. (**G-I**) Immunocytochemistry of pluripotency markers OCT4 (G), NANOG (H) and TRA-1-60 (I) in *cPEX5^iKO^*; nuclei were counterstained with Hoechst 33342 (blue); scale: 100 µm. (**J**) Molecular karyotyping using a genome-wide SNP array demonstrated chromosomal stability of hiPSCs after genome editing. (**K**) Flow cytometric analysis of cardiac troponin T (cTNT) stained hiPSC-CMs was performed at day 60 post-cardiac differentiation. Original measurement from *cPEX5^iKO^*shows 88.6 % of cTNT^+^ cells obtained after cardiomyocyte selection. (**L**) Western blot confirms PEX5 protein deficiency in *cPEX5^iKO^*. For uncropped images see Figure S9. (**M,N**) Staining of the membrane protein Pex14 shows decreased number and increased size of peroxisomes in *cPEX5^iKO^*. Scale: 10 µm. (N) Quantification of (M). Peroxisome number normalized to nuclei. N = 16 (WT) and 25 (*cPEX5^iKO^*). (**O**) Co-staining of α-Actinin with ACAA1 shows successful generation of hiPSC-CMs with loss of ACAA1 in *cPEX5^iKO^*; scale: 10 µm. (**P-R**) Co-staining of PMP70 with peroxisomal matrix proteins ACAA1 (P), HSD17β4 (Q) or ACOX1 (R) shows peroxisomal import defect; scale: 10 µm.

Next, hiPSCs were differentiated into spontaneously beating ventricular-like cardiomyocytes (hiPSC-CMs). After differentiation, 83 ± 3 % (n=15) of the cells were positive for cardiac troponin T (cTNT) (Figure 1L and S1A). We denoted the knock-out cardiomyocytes as c*PEX5^iKO^*. Western blotting proved absence of PEX5 in *cPEX5^iKO^* (Figure 1M and S9). As the antibody does not recognize the truncated protein, we confirmed that a potential residual protein fragment is not functional. Staining the peroxisomal membrane protein (PMP) PEX14 in *cPEX5^iKO^* showed a decreased number of peroxisome structures with significantly increased size (Figure 1N,O and S1B). We analyzed the residual peroxisomal structures for the presence of peroxisomal matrix proteins. All tested marker proteins, 3-ketoacyl-CoA thiolase (acetyl-Coenzyme A acyltransferase 1, ACAA1) (Figure 1P,Q and S1C,D), peroxisomal multifunctional enzyme type 2 (‘hydroxysteroid 17β-dehydrogenase 4’, HSD17β4) (Figure 1R and S1F), and acyl-CoA oxidase 1 (ACOX1) (Figure 1S and S1E) were absent from peroxisomes. Unimported HSD17β4 is likely unstable in the cytosol as shown by Western blotting (Figure S1G and S9). The enlarged, virtually empty and non-functional membrane organelles with reduced abundance represent the typical PBD phenotype. Such matrix protein-free structures that can only be recognized by the presence of peroxisomal membrane proteins, have been previously designated as peroxisomal ‘ghosts’ and have been observed in otherwise peroxisome-rich tissues of PBD patients and in mouse models^10^. Taken together, we established a peroxisomal protein import defect in c*PEX5^iKO^*. Peroxisomal matrix proteins were absent from the peroxisomal ghosts and mostly degraded in the cytoplasm.

### Peroxisome-deficiency linked to mitochondrial dysfunction

Peroxisomal dysfunction is often accompanied by mitochondrial structural defects^11^ and peroxisomes act in concert with mitochondria to provide energy homeostasis^12^. As the heart is a metabolically demanding organ and highly depending on mitochondrial energetics, we interrogated if mitochondrial functionality is preserved in the *cPEX5^iKO^*. In order to analyze whether mitochondrial structure is affected by *PEX5* deficiency, we stained these organelles in hiPSC-CMs with MitoTracker green. By structured illumination microscopy (SIM) (Figure 2A) we observed an increased number of mitochondria per cell (Figure 2B). These mitochondria show a decreased number of branches per mitochondria (Figure 2C) with decreased branch length (Figure 2D) together with a decreased number of branch junctions (Figure 2E) indicating massive mitochondrial fragmentation in *cPEX5^iKO^* cells. To assess whether this altered morphology is associated with impaired function, we measured mitochondrial membrane potential and found a significant reduction compared to wild-type (WT) (Figure 2F). Both, fragmentation and reduced membrane potential, suggest mitochondrial dysfunction with defects in oxidative phosphorylation and ATP production. In respirometry measurements (Figure 2G), basal oxygen consumption rate (OCR) and ATP-linked OCR were not altered (Figure 2H,I), but maximal respiration and spare respiratory capacity in *cPEX5^iKO^* cells were significantly decreased (Figure 2J,K). Beating frequency, the total DNA content, the DNA to cell conversion or the number of cells per well were not different between WT and *cPEX5^iKO^* (Figure S2A-D). In whole cell lysates, we found decreased ATP levels (Figure 2L), implying that peroxisome-induced mitochondrial dysfunction affects cellular energy supply.

**Figure 2:**
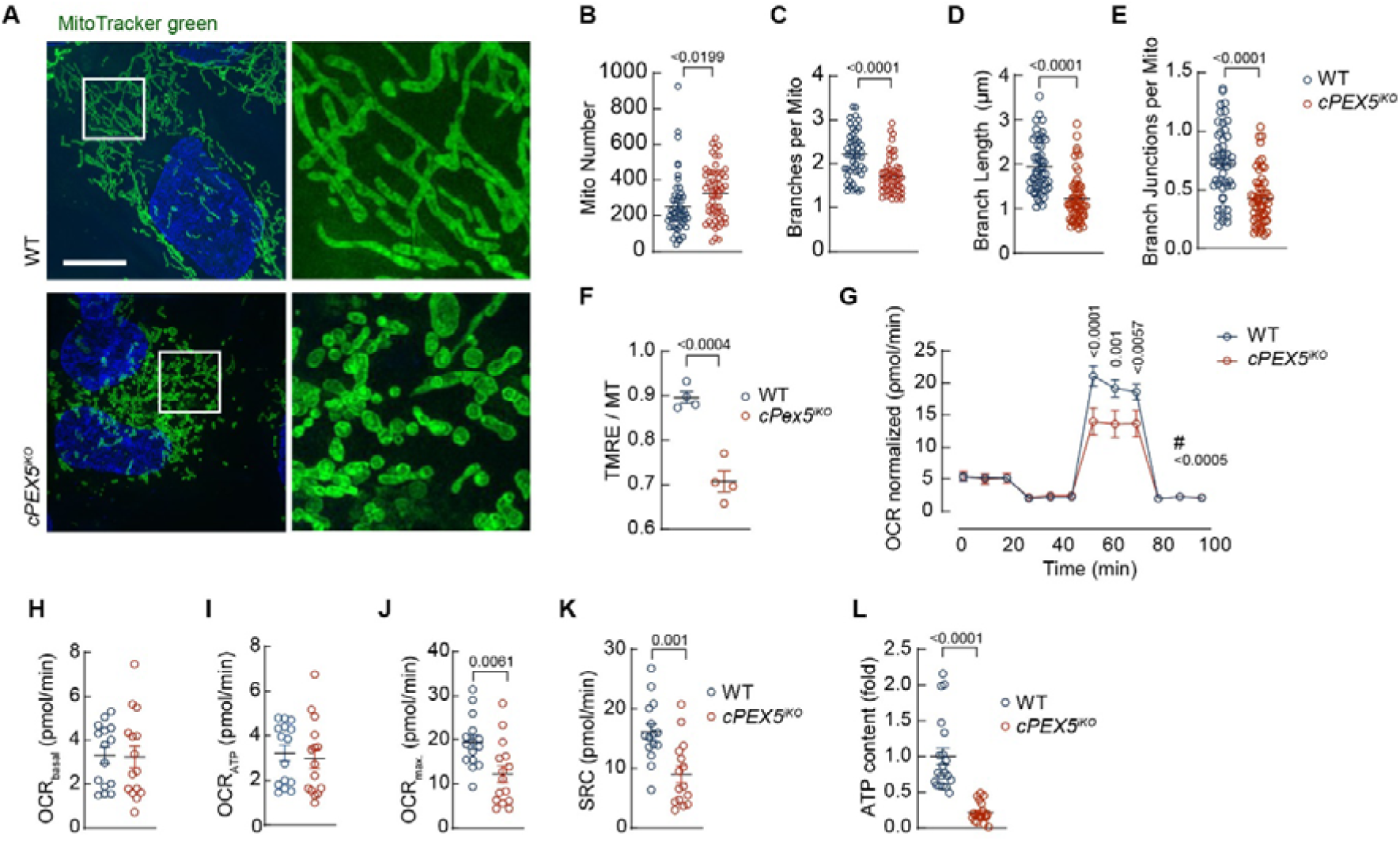
Mitochondrial fragmentation and defective oxygen-consumption result in reduced ATP availability in the peroxisome-deficient cardiomyocytes. **(A)** Mitochondrial fragmentation in *cPEX5^iKO^* iPSC-CMs observed by Mitotracker green staining. Structured illumination microscopy; Scale: 10 µm (**B-E**) Quantification of (A); Mitochondria analyzer ImageJ. Increased number of mitochondria per cell (B) with reduced number auf branches (C), decreased branch length (D) and decreased branch junctions per mitochondria (E). (**F**) TMRE (tetramethylrhodamin, 600nM) measurement shows reduced mitochondrial membrane potential in *cPEX5^iKO^*. Normalized to MitoTracker green (MT). (**G-K**) Measurement of oxygen consumption rate (OCR, two-way ANOVA) in hiPSC-CMs (G) revealed no change in basal respiration (H) or ATP-linked OCR (I), but significant deficits in maximal respiration (J) and spare respiratory capacity (K) in *cPEX5^iKO^* compared to WT cells. Normalized to cell number (nuclei). N=15 per group, Welch’s test. (**L**) ATP Assay in whole cell lysates. Normalized to WT, N=20 per group.

### Peroxisome-deficient engineered human myocardium (EHM) presents with heart failure phenotype

To analyze the functional impact of *PEX5* deficiency and its consequences on cardiac contraction and force generation, we decided to develop a human cardiac muscle model (engineered human myocardium, EHM) from WT and *cPEX5^iKO^* cardiomyocytes (Figure 3A and Videos S1/S2). In contrast to the more immature state of hiPSC-CMs, these three-dimensional cardiac model systems show improved recapitulation of functional and morphological features of the human heart^13,14,14^, so that force production and electromechanical coupling of cardiomyocytes can be assessed in a functional syncytium. WT and *cPEX5^iKO^* were mixed with WT human fibroblasts in a 70:30 ratio in a collagen matrix and seeded into molds of a 48-well EHM plate^15^. This culture plate allows the formation of 48 EHM in parallel around flexible posts that confer mechanical loading and are utilized for optical analysis of EHM function by automated tracking of pole movement. WT and *cPEX5^iKO^*EHM were generated and analyzed up to 43 days in culture. Both, WT and *cPEX5^iKO^* cardiomyocytes formed EHM with comparable macroscopic appearance (Figure 3A). Cardiomyocytes from *cPEX5^iKO^* EHM show the peroxisomal ghost phenotype with reduced number and increased size compared to peroxisomes from WT EHM cardiomyocytes, whereas, as expected, the phenotype of the fibroblasts was not altered in the WT or *cPEX5^iKO^* EHM (Figure 3B-F). Quantification revealed that *cPEX5^iKO^* EHM contained less and smaller cardiomyocytes (Figure 3G, H). In accordance with the observed mitochondrial energy deficit, optical imaging of *cPEX5^iKO^* EHM function revealed that EHM shortening was markedly reduced as early as day 14 of EHM development (Figure S3A). Force measurements at increasing Ca^2+^ concentrations under isometric load in thermostat-equipped organ baths at day 43 of culture confirmed a significantly decreased force of contraction (Figure 3I). Resting force as an indicator of passive tissue tension was increased in *cPEX5^iKO^* EHM (Figure 3J) and active to passive force ratio was significantly decreased (Figure 3K), implicating a HF phenotype^16^. When challenged by an increase in frequency, we observed a reduction of force in *cPEX5^iKO^*EHM (i.e., negative force-frequency relationship, also known as Bowditch Treppe^17^) but not WT EHM (Figure 3L). Deficits in frequency-dependent force potentiation are amongst the most important functional changes in HF with reduced ejection fraction^18,19^. When normalized to cardiomyocyte number, force generation of *cPEX5^iKO^* EHM remained significantly reduced (Figure S3B). In summary, loss of functional peroxisomes disturbs force generation in the EHM, identifying peroxisomes as substantial contributors to heart muscle function.

**Figure 3:**
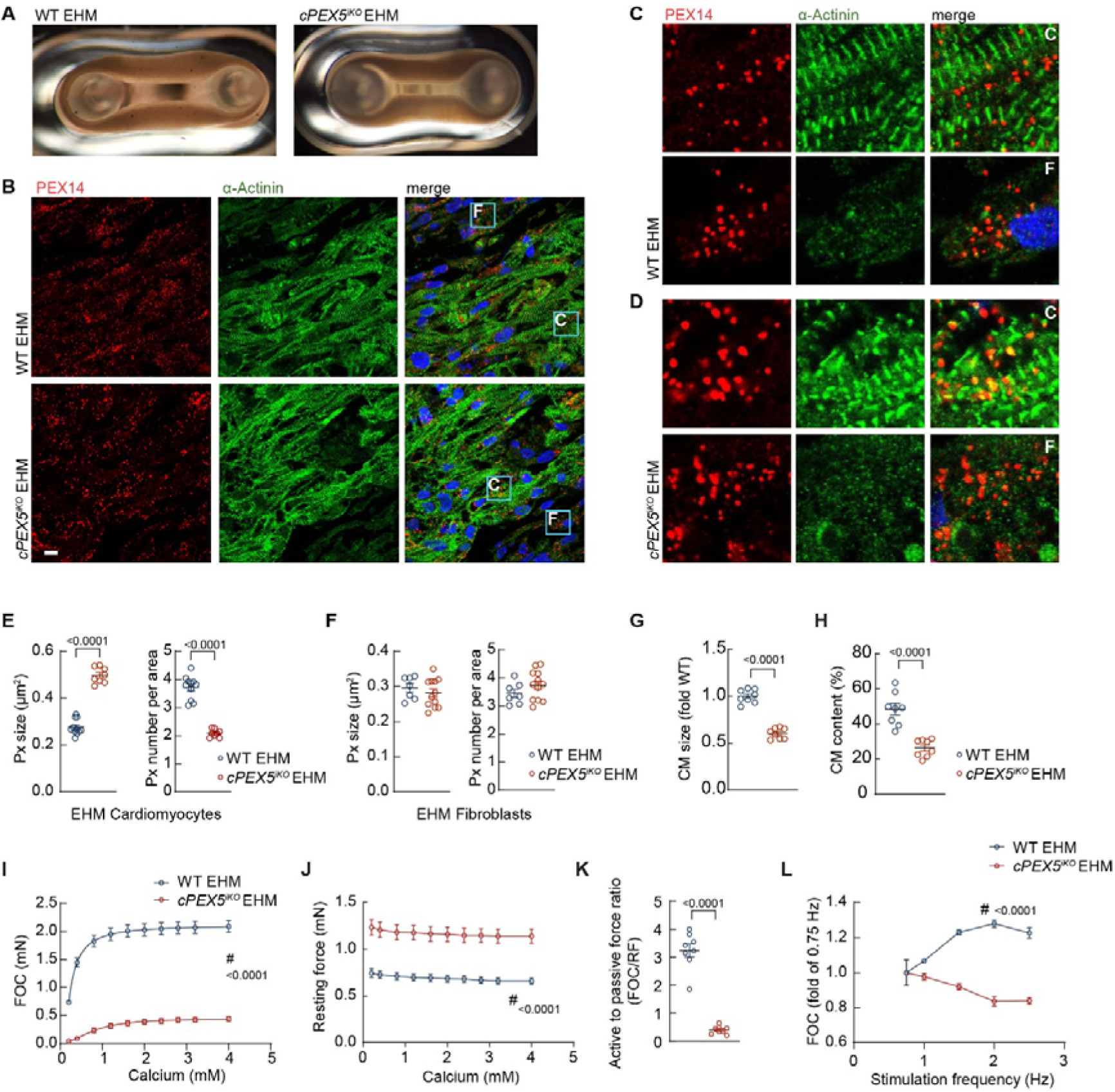
Heart failure phenotype in *PEX5*-deficient engineered human myocardium. **(A)** EHM were generated from human WT and *cPEX5^iKO^* iPSC-CMs. See Video S1. (**B-F**) Verification of peroxisomal biogenesis deficiency in EHM. PEX14 staining of peroxisomes in EHM shows increased size and reduced number of peroxisomes in *cPEX5^iKO^* EHM cardiomyocytes (D,E), but no change in peroxisome size or number in fibroblasts of *cPEX5^iKO^* EHM (D,F). WT EHM peroxisomes did not change (C,E,F). N=11 WT and 8 *cPEX5^iKO^* EHM. Scale: 10 µm. F = fibroblasts, C = cardiomyocytes. (**G**) Cardiomyocyte size (FSC-A median) and (**H**) cardiomyocyte content were reduced in *cPEX5^iKO^* EHM. N=8 EHM per group. ^#^two-way ANOVA; Bonferroni’s multiple comparisons. (**I-L**) Organ bath experiments reveal (I) lower force of contraction (FOC), (J) increased resting force, and (K) pathological (<1) active to passive force ratio in *cPEX5^iKO^* EHM. *cPEX5^iKO^* EHM also show an impaired (negative) force frequency response (L). N=8 EHM per group. ^#^two-way ANOVA. Px = peroxisome. See also Videos S1 and S2.

### Altered Ca^2+^ handling and irregular conduction upon peroxisome-induced mitochondrial dysfunction result in heart failure phenotype

Altered Ca^2+^ handling is one of the hallmarks of HF. Hence, we determined cytosolic Ca^2+^ concentrations and their post-rest potentiation in EHM as an indicator for sarcoplasmic reticulum (SR) Ca^2+^ load. After pausing electrical stimulation for 10 seconds in *cPEX5^iKO^* EHM, post-rest potentiation was significantly reduced compared to WT (Figure 4A,B), indicating impaired SR Ca^2+^ loading. Therefore, we analyzed hiPSC-CMs after loading the cells with the cytosolic Ca^2+^ indicator dye Fluo-4 AM. Although spark frequency was not increased and a Ca^2+^ leak was not detected in *cPEX5^iKO^* cardiomyocytes (Figure S4A,B), we observed Ca^2+^ transients with significantly prolonged time-to-peak (Figure 4C,D) and prolonged Ca^2+^ decay time (Figure 4C,E) in *cPEX5^iKO^*, with slight (non-significantly) reduction in the Ca^2+^ transient amplitude (Figure 4C,F). Together, the slower decay of the Ca^2+^ transients together with reduced post-rest potentiation indicate that reduced SERCA activity and SR Ca^2+^ load contribute to reduced force generation. The phenotypes identified in these experiments point to an essential role of peroxisomes in the main ATP-dependent processes involved in muscle contraction: cross-bridge cycling and the maintenance of Ca^2+^-homoeostasis.

**Figure 4:**
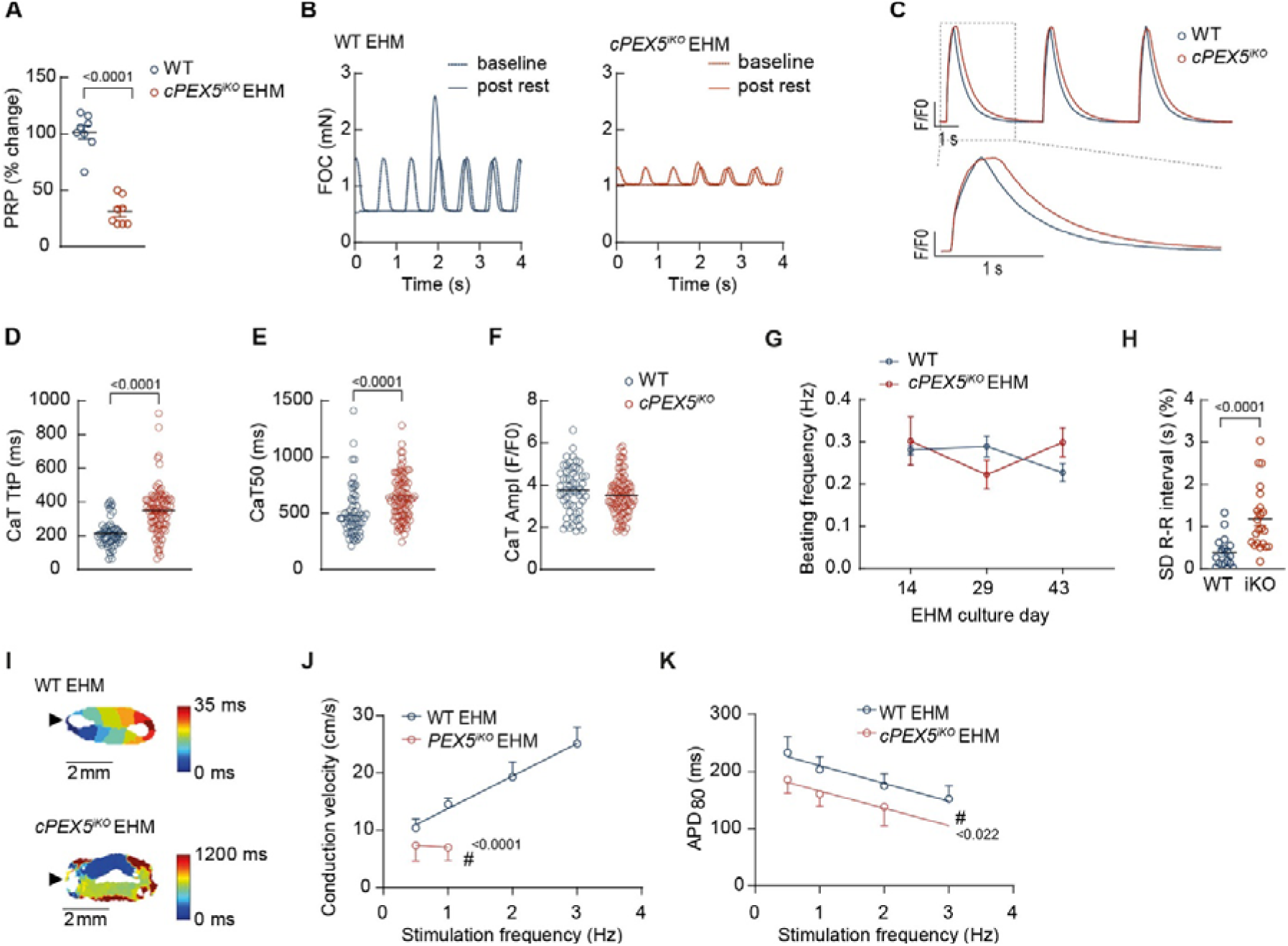
Defects in Ca^2+^-handling and electrical conduction upon *PEX5* deficiency explain heart failure phenotype. (**A,B**) Ca^2+^-handling in EHM. Post-rest potentiation (PRP) of force (force of contraction, FOC) after pausing electrical stimulation for 10 s is decreased in *cPEX5^iKO^* EHM. (B) Representative traces of (A). (**C-F**) Ca^2+^ measurement in hiPSC-CMs. (C) Original recordings of Ca^2+^ measurements. No change in Ca^2+^ transient amplitude (CaT Ampl, D), but prolonged time to peak (TtP, E) and relaxation time (CaT50, F). N = 58-92 cells per group. Mann-Whitney test. (**G,H**) EHM reveal comparable beating frequency (G) but increased beat to beat variability (H). SD [standard deviation] of beat-to-beat interval [R-R] at day 43, Welch’s test (H). (**I-K**) Activation map (I) of WT and *cPEX5^iKO^* EHM with colors representing the time point of maximal signal amplitude after electrical point stimulation (arrowhead). (J) Conduction velocity. (K) Frequency dependence of action potential duration at 80% repolarization (APD_80_). N =3-4 EHM, two-way ANOVA. See also Videos S3 and S4.

Reduced SR Ca^2+^ load may also contribute to arrhythmia. Interestingly, the spontaneous beating rate of the EHM was comparable between WT and *cPEX5^iKO^* EHM (Figure 4G). However, we observed an increased beat-to-beat variability in *cPEX5^iKO^*EHM (Figure 4H). Therefore, we used optical mapping of WT and *cPEX5*^iKO^ EHM to follow electrical conduction in these model systems. WT EHM (4 out of 4) showed homogeneous excitation and frequency-dependent action potential duration (APD) and conduction velocity (Figure 4I-K and Video S3). The mean conduction velocity was 19.5 cm/s at 2 Hz. The WT vector map shows the excitation wave origin and proper wave propagation and conduction velocity increases with an increase in pacing frequency in 3 of the 4 tissues (Figure 4I,J). In contrast to that, all (4 out of 4) *cPEX5^iKO^* showed incoherent excitation in multiple areas with slow and incomplete conduction (Figure 4I, K and Video S4). Also, APD was significantly decreased in *cPEX5^iKO^* EHM (Figure 4I,K). This drastically aberrant and largely disrupted propagation of the electrical stimulation supports the notion that *PEX5* deficiency leads to arrhythmic vulnerability on tissue level, that in a more physiological scenario would lead to HF.

In summary, in the engineered heart model for peroxisome biogenesis deficiency, the absence of functional peroxisomes induces mitochondrial dysfunction in association with aberrant SERCA function, an SR Ca^2+^ leak, reduced force generation and defective conduction. On tissue level, the irregular excitation propagation ultimately results in a HF-like phenotype.

### A mouse model for peroxisome biogenesis deficiency in the heart

To substantiate these findings, we decided to create and analyze an orthogonal murine model. We generated a cardiomyocyte-specific knockout of the *Pex5* gene in mice (Myh6-Cre^+/-^ Pex5^fl/fl^, Pex5^cKO^). In this model, the cardiomyocyte-specific α-myosin heavy chain (*Myh6*)^20^ promotor drives the deletion of exons 11 to 14 of the *Pex5* gene^21,22^ mediated by Cre recombinase (Figure 5A). Cre-negative, homozygous floxed Pex5 mice (Myh6-Cre^-/-^ Pex5^flox/flox^, WT) and WT Pex5 mice expressing Cre recombinase (Myh6-Cre^+/-^ Pex5^wt/wt^, WT^c^) were used as control groups. We verified the heart-specific knockout of *Pex5* by PCR (Figure S5A). Homozygous Pex5^cKO^ mice are viable and are healthy at birth.

**Figure 5:**
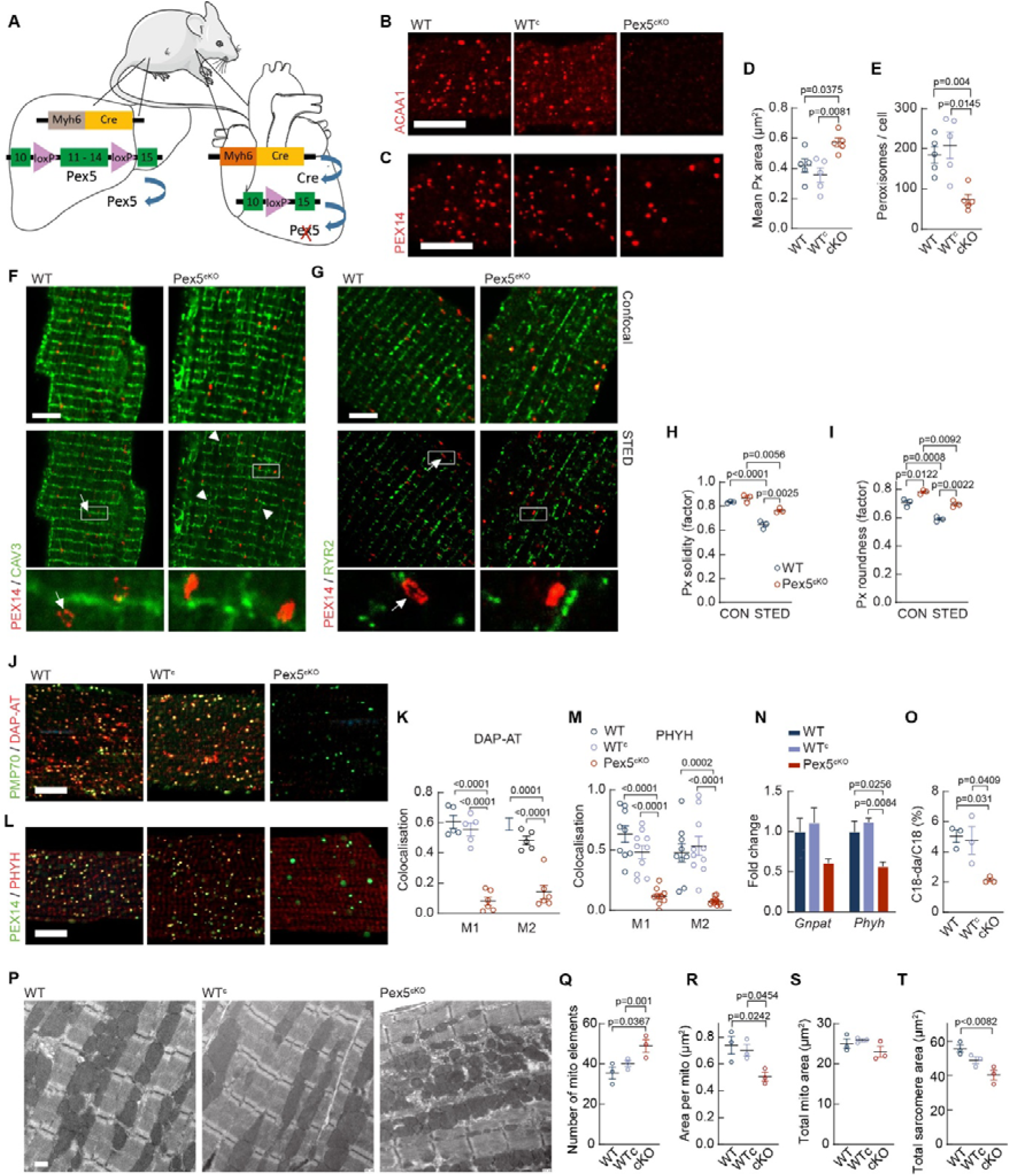
Peroxisomal enzyme deficiency in the murine heart leads to mitochondrial fragmentation. **(A)** Cre-driven cardiac-specific *Pex5* knockout (Pex5^cKO^). (**B,C**) IF of isolated cardiomyocytes shows peroxisomal ghosts (B, anti-PEX14) with increased area and deceased number of peroxisomes per cell and loss of ACAA1 (C) in peroxisomes. Scale: 10 µm. (**D**) Quantification of PEX14 staining (B). N = 30 cells, one-way ANOVA, Tukey’s multiple comparisons. (**F,G**) Confocal (upper row) and STED (lower row) microscope imaging of isolated cardiomyocytes from Pex5^cKO^ mice stained for PEX14 and CAV3 (F) or RYR2 (G) indicates increase in axial T-tubule structures (arrowheads). Arrows indicate peroxisomal lumen in WT cardiomyocytes, which is not seen in Pex5^cKO^ cardiomyocytes. Scale: 5 µm. (**H,I**) Analysis of peroxisomes in confocal and STED images (F,G). Solidity of peroxisomes calculated as ratio of the area and convex area reveals that peroxisomes in Pex5^cKO^ are more solid (H). This can be seen with STED, but not confocal microscopy. Roundness of peroxisomes is calculated based on the area and major axis. Peroxisomes in Pex5^cKO^ are more round compared to WT (I). STED is a more sensitive approach than confocal when imaging less round objects. N = 3 mice, one-way ANOVA, Tukey’s multiple comparisons. (**J-M**) Loss of DAP-AT (J) and PHYH (L) in peroxisomes of isolated cardiomyocytes of Pex5^cKO^ mice. Scale: 10 µm. (K,M) Quantification of J,L (Manders coefficient). M1= Fraction of PEX14 / PMP70 that overlaps with PHYH or DAP-AT, M2= Fraction of PHYH or DAP-AT that overlaps with PEX14/PMP70. N = 5 - 9 cells per group, one-way ANOVA, Tukey’s multiple comparisons. (**N**) Reduction of *Gnpat* and significant downregulation of *Phyh* mRNA. qPCR. One-way ANOVA, Tukey’s multiple comparisons. N=3 mice per group. Normalized to *Gapdh*. (**O**) Loss of C18 plasmalogens in Pex5^cKO^ hearts. N = 3 hearts per group, one-way ANOVA, Tukey’s multiple comparisons. (**P**) EM images of heart tissue sections show mitochondrial fragmentation and sarcomere loss in Pex5^cKO^ hearts. Scale: 500 nm. (**Q-T**) Quantification of (P). N = 3 mice per group, one-way ANOVA, Tukey’s multiple comparisons.

Knockout of *Pex5* led to deficient import of PMP, as confirmed in isolated cardiomyocytes. Immunofluorescence detection of ACAA1, a protein whose import is dependent on PEX5 and its co-receptor PEX7^23^, revealed that ACAA1 is apparently absent in Pex5^cKO^ cells, likely because the non-imported protein is unstable in the cytosol (Figure 5B). Staining of PEX14 shows reduction of peroxisome number with significant increase in apparent peroxisome size (Figure 5C-E), confirming that peroxisomal structures are present in Pex5^cKO^ as empty and non-functional ‘ghosts’. Levels of PMP70, however, were unaltered in the Pex5^cKO^ hearts, in line with the expression in peroxisomal ghosts (Figure S5B).

We analyzed WT and Pex5^cKO^ peroxisomal morphology by STED microscopy (Figure 5F,G) revealing finer details of peroxisomal substructure^9^. In WT cardiomyocytes, the lumen of the peroxisomes was readily detectable. In Pex5^cKO^, ghosts appeared compacted without detectable lumen and morphology was drastically altered (Figure 5F,G). We quantified the roundness and solidity of the peroxisomes as shape parameters in confocal and STED microscopy images and found Pex5^cKO^ peroxisomes to be more round and solid compared to control (Figure 5H, I and S5C).

To assess the functional changes in Pex5^cKO^ mice, we examined enzymes involved in plasmalogen synthesis and α-oxidation of fatty acids, two fundamental metabolic functions of peroxisomes with a potential specific role in the heart^24^. In the healthy heart, about 25% of lipids are plasmalogens^25^ and accumulation of the α-oxidation substrate phytanic acid is cardiotoxic^26^. Hence, we isolated cardiomyocytes from WT and Pex5^cKO^ mice to investigate expression of the peroxisomal enzymes Dihydroxyacetone phosphate acyltransferase (DAP-AT, encoded by *Gnpat*, Figure 5J) and Phytanoyl-CoA dioxygenase (Phytanoyl-CoA hydroxylase, PHYH, Figure 5L), involved in plasmalogen synthesis and α-oxidation, respectively. Both enzymes were undetectable in Pex5^cKO^ cardiomyocytes (Figure 5J-M and S5D,E). On the mRNA level, *Phyh* was downregulated in Pex5^cKO^ hearts, and *Gnpat* was not significantly changed (Figure 3N). Blood plasmalogen levels were unaltered (Figure S5F) and C16 plasmalogens were non-significantly reduced (Figure S5G), but C18 plasmalogens were significantly decreased in Pex5^cKO^ hearts (Figure 3O). These findings show a heart-specific loss of plasmalogens in Pex5^cKO^ mice. It can be concluded that these are autonomously produced in the heart tissue and not directly obtained through circulation.

### Peroxisome biogenesis deficiency induces mitochondrial fragmentation in the murine heart

Plasmalogen loss is known to link peroxisome to mitochondrial dysfunction^27^. Notably, electron microscopy of Pex5^cKO^ heart tissue sections showed an increase in number (Figure 5P,Q) and a reduction in size (Figure 5P,R) of the Pex5^cKO^ mitochondria, whereas total mitochondrial area remained unchanged (Figure 5P,S). Hence, analogous to the human *Pex5^iKO^* cells, peroxisomal dysfunction results in mitochondrial fragmentation in the murine cardiomyocytes.

Likewise, we observed cardiomyocyte degeneration, sarcomere loss and accumulation of vesicles and electron dense material in these sections (Figure 5P,T). To assess a potential change in mitochondrial protein content, we isolated mitochondria from cardiac tissue and analyzed mitochondrial protein expression. Carnitine palmitoyltransferase 1B (CPT1B, fatty acid metabolism), Cytochrome c oxidase subunit 5A (COX5a), Cytochrome c oxidase subunit 4I1 (COX-4I1, both complex IV) and ATP synthase F1 subunit beta (ATP5B, complex V) were unchanged in relation to the stable Voltage-dependent anion-selective channel protein (VDAC) in the outer mitochondrial membrane (Figure S5H,I). We also tested respiratory chain assembly using native gel electrophoresis. Analysis of complex II with Succinate dehydrogenase complex, subunit A (SDHA) and respiratory chain supercomplexes with antibodies directed against Rieske (complex III) and COX5a (complex IV) revealed no compositional changes of the respiratory chain complexes (Figure S5J,K). Hence, increased mitochondrial fragmentation does not affect the expression of the main mitochondrial proteins or structural assembly of the respiratory chain complexes. This comparably mild mitochondrial phenotype is compatible with the apparently healthy birth of the *Pex5^iKO^* mice and a potential developmental phenotype.

### Dilated cardiomyopathy and heart failure caused by peroxisome import deficiency in the murine heart

Although Pex5-deficient mice appear healthy at birth and during their first weeks of life, we observed a high mortality of Pex5^cKO^ mice starting with about four months of life, with a mean survival of 150 days (Figure 6A and S6A). Body weight of the animals was monitored from birth until 23 weeks of age. Throughout this period, no significant differences were observed between the Pex5^cKO^ mice and WT controls (Figure S6B). Cardiac function and geometry of Pex5^cKO^ and WT mice were assessed through serial echocardiographic evaluations conducted between 7 and 23 weeks of age (Figure 6B-K and S6C-O). At 7 and 10 weeks, no significant differences were observed between WT and Pex5^cKO^ mice in terms of heart rate, left ventricular mass (LVM)/BW and LVM (Figure 6B,C and S6C). However, LVM/BW ratio and LVM of Pex5^cKO^ mice were increased versus WT mice at the later time points (Figure 6C and S6C). The LV internal diameters in end diastole and in end systole (LVIDd, LVIDs) were enlarged at a constant level between week 7 and 15, and progressively increased further from week 15 (Figure 6D,E,F and S6D). Additionally, we found signs of dilation of the entire heart in the Pex5^cKO^ mice with constant enlargement of systolic and diastolic circumference and heart volume from week 7 to 15, followed by a progressive increase of these parameters (Figure S6E-I). The anterior wall thickness in diastole and systole was slightly decreased, wall thickening fraction was reduced from week 15 onwards (Figure 6G and S6J,K). The posterior wall thickness decreased in systole but not in diastole (Figure S6L,M). The relative wall thickness, a measure for geometry alterations of the LV, was constantly decreased from 7 to 23 weeks (Figure 6H). In line with these structural deficiencies, we found a progressive decline in functional parameters. The ejection fraction, the percentage of blood that is pumped out by the LV during one heartbeat, and the contraction of the ventricle were significantly decreased from week 7, and worsened over the time course from week 7 to 23 (Figure 6I and S6N, O). Finally, we also looked at radial diastolic peak velocity and reverse longitudinal strain rate as markers for the diastolic function of the heart. Already with 10 weeks of age, these were significantly decreased in the Pex5^cKO^ (Figure 6J,K). Hence, the echocardiographic data show that Pex5^cKO^ mice develop a severe DCM with significant decline in diastolic function resulting in HF and mimicking the *cPEX5^iKO^*EHM phenotype.

**Figure 6:**
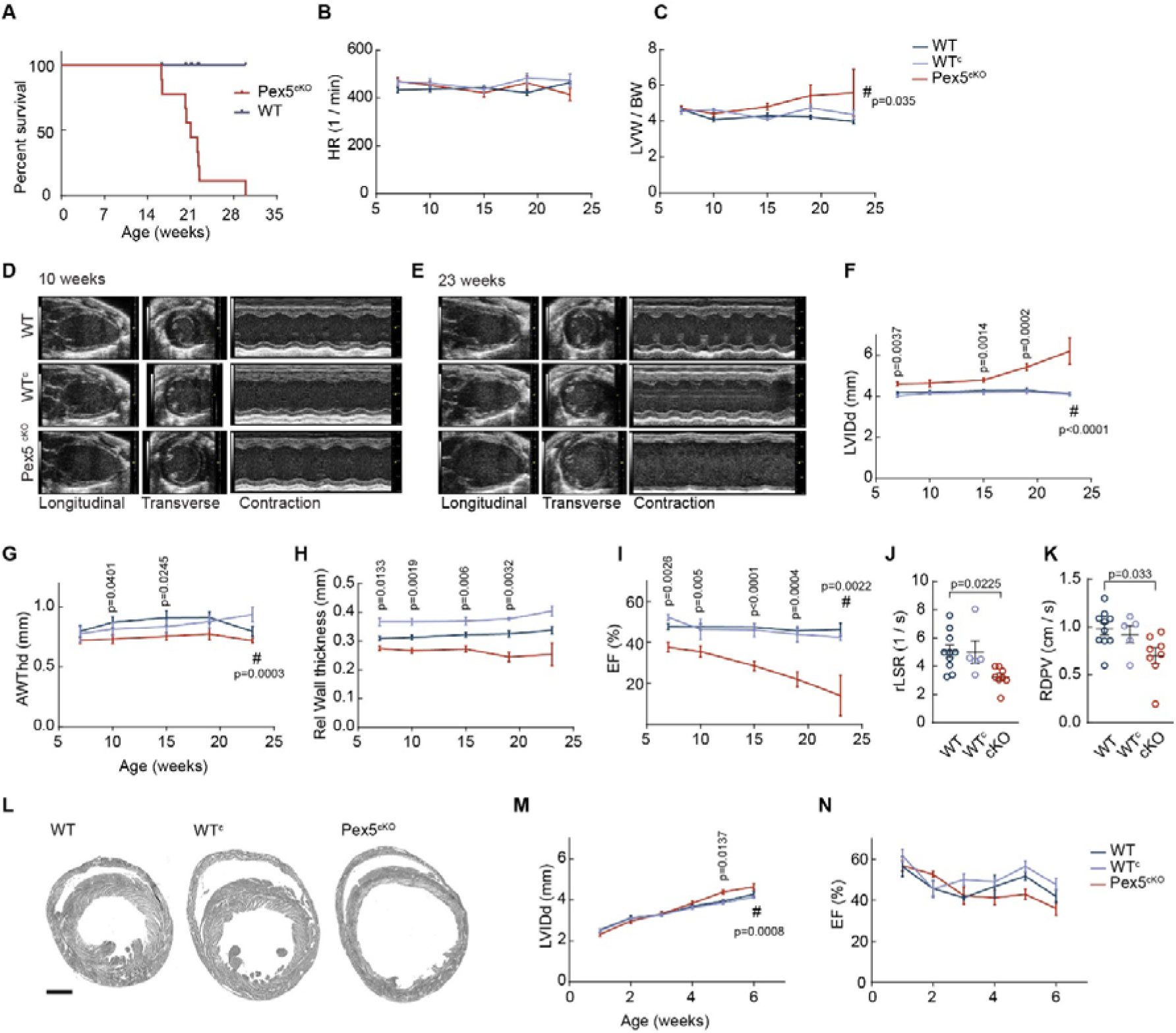
Dilated cardiomyopathy and heart failure due to cardiac *Pex5* deficiency. **(A)** Kaplan Meier survival curve shows early death of Pex5^cKO^ mice (mean survival: 150 days). N = 11 mice per group. (**B-K**) Echocardiography of 7- to 23-week-old mice shows dilated hearts with decreased contraction indicating severe DCM in Pex5^cKO^ mice. Function decreases over time. N = 3-16 mice per time point per group. Two-way ANOVA, Tukey’s multiple comparisons. (B) HR = heart rate, (C) LVM/BW ratio; BW = body weight, LVM = left ventricular mass. (D) representative images of transverse and longitudinal recordings and contraction profile in 10-(D) and 23-week (E) old mice. (F) LVIDd = left ventricular internal diameter in diastole, (G) AWThd = anterior wall thickness in diastole, (H) relative wall thickness (two times posterior wall thickness divided by LVIDd), (I) EF = ejection fraction, (J,K) Echocardiography of 10-week-old mice shows significant decrease in parameters for diastolic function. rLSR: reverse longitudinal strain rate and RDPV: radial diastolic peak velocity. N = 5-10 mice per group, one-way ANOVA, Tukey’s multiple comparisons. (**L**) HE-stained cross-sections show enlarged ventricles with decreased wall thickness in Pex5^cKO^ hearts (age = 22 weeks). Scale 1 mm. (**M,N**) Echocardiography of 1- to 6-week-old mice shows that first significant signs of cardiac dilation are observed in 5-week-old mice. N = 5-7 mice per time point per group. Two-way ANOVA, Tukey’s multiple comparisons.

We prepared whole heart cross-sections of 22-week-old mice, stained them with hematoxylin-eosin (HE) and reconstructed the optical slices by assembling the individual images. The data confirm the impressive dilation phenotype with decreased wall thickness (Figure 6L). Histological analysis of HE-stained heart sections revealed tissue damage in the apparent absence of fibrosis (Figure S6P). CD68 staining showed macrophage infiltration in Pex5^cKO^ hearts (Figure S6P). HE-stained lung sections of Pex5^cKO^ mice did not show signs of pathology (Figure S6Q).

Next, we asked whether this phenotype was present earlier in life and performed echocardiography weekly after early genotyping (at one week of age) until six weeks of age. Heart rate, LVM or LVM/BW ratio were not changed (Figure S6R). At the earliest time point (weeks 1-3) we also found no effect on heart size, as LVID, circumference and heart volume (Figure 6M and S6S) did not change. However, from week 4 onwards, these parameters increased with first significant differences observed in week 5 (Figure 6M and S6S). Taken together, these data show that development of cardiac dilation starts at 4 weeks of age without an overt effect on LV geometry or heart function at this age (Figure 6N and S6T). Hence, the Pex5^cKO^ phenotype is initially asymptomatic and HF pathology develops gradually after weaning, resulting in early death at an age of about five months. From a broader perspective, our data show that cardiac peroxisome deficiency is associated with strongly interconnected metabolic and developmental phenotypes.

Mice expressing Cre recombinase, Myh6-Cre^+/-^ Pex5^wt/wt^ (WT^c^), developed a mild cardiac phenotype at about three to six months of age, that has been described before^28,29^, confirming the importance of using these mice as a second control group together with Myh6-Cre^-/-^ Pex5^flox/flox^ (WT) mice. Survival of WT^c^ mice is not affected (Figure S6A). In contrast to the dilated heart phenotype observed in Pex5^cKO^ mice, WT^c^ mice show increase in wall thickness (Figure 4H and S6J,M), but no change in the functional parameters (Figure 6I and S6N,O). Histological characterization of WT^c^ hearts did not show signs of fibrosis, whereas inflammation and limited background level of macrophage infiltration could be detected in WT^c^ hearts, albeit at a far lower level than in Pex5^cKO^ (Figure S6P). This phenotype is likely caused by Cre expression^28,29^.

### Peroxisome deficiency impairs electrical conduction

The murine model developed here shows that peroxisome deficiency causes a severe cardiac phenotype that mirrors the phenotype observed in the human model. Next, we wanted to study how these deficits translate to the electrophysiology of the whole heart. We used Langendorff-perfused hearts to record bipolar electrograms in explanted hearts from Pex5^cKO^ mice to determine PR, QRS, QT-Intervals and the effective refractory period (ERP) (Figure 7A-D and S7A). Despite no change in the spontaneous beating rate (Figure S7B), the QRS duration (indicating the conduction velocity of the electrical activation within the ventricles) was markedly prolonged in Pex5^cKO^ hearts (Figure 7B). Similarly, the QT interval (indicative of the average duration of ventricular action potentials) was significantly prolonged in Pex5^cKO^ mice (Figure 7C). Furthermore, the ERP (a measure of the minimum time interval between successive action potential excitation) was substantially increased in Pex5^cKO^ hearts (Figure 7D). These electrical alterations are typical for failing hearts and are known to be pro-arrhythmic and thus may account for the sudden cardiac death that we observe in the peroxisome-deficient mice. In contrast to the prolonged QRS and QT intervals, our ex-vivo Langendorff experiments revealed a shortening of the PR interval in Pex5^cKO^ mice compared to WT controls (Figure S7G). This finding suggests an accelerated atrial conduction velocity or atrioventricular conduction in the absence of PEX5. This unexpected finding highlights the complex and potentially tissue-specific impacts of PEX5 deficiency on cardiac electrophysiology, warranting further investigations into cardiac conduction and ion channel function.

**Figure 7:**
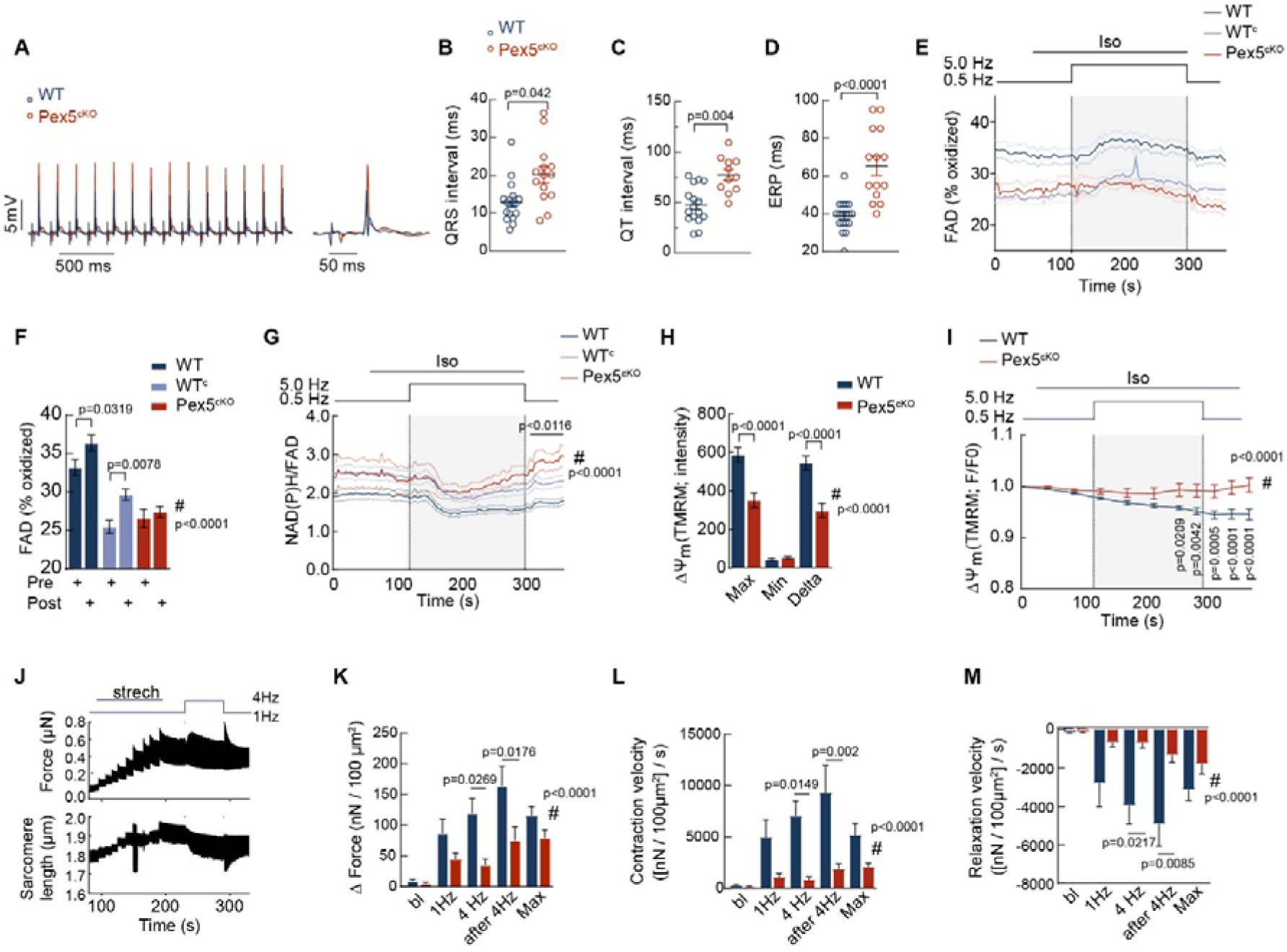
Defects in mitochondrial membrane potential and energetics result in defective electrical conduction, decreased cellular contractility and reduced SR Ca^2+^storage in Pex5^cKO^ myocytes. (**A-D**) Electrophysiological characterization of cardiac electrograms. (**A**) Original cardiac electrogram (ECG) recording of explanted hearts from a WT heart (black) and a Pex5^cKO^ heart (red). (B-D) Statistical analysis of ECG parameters during spontaneous beating. QRS duration, 12.87 ± 0.12 vs. 20.20 ± 0.22 ms (B), QT interval, 48.2 ± 0.4 ms vs. 73.6 ms ± 0.5 ms (C) and ventricular effective refractory period (ERP), 38.9 ± 1.9 vs. 65.4 ± 5.0 ms (D) in WT and Pex5^cKO^ hearts. N = 14 - 18 hearts per group. (**E-G**) Autofluorescence of NAD(P)H and FAD were measured in murine cardiomyocytes exposed to the indicated protocol (top). Level of oxidized FAD is reduced compared to WT and did not show an increase following workload transition (E, F); Pre = basal FAD level at 0,5 Hz stimulation; Post = 40-60 s after initiation of 5 Hz stimulation. (G) NADH oxidation following workload transition is slightly reduced in cKO compared to WT cells and the NAD(P)H redox state after the initial oxidation cannot be maintained throughout the stress protocol in Pex5^cKO^ compared to WT cells. N = 6-21 cells from 3 mice per group. (**H, I**) Cardiomyocytes were loaded with TMRM to measure changes in the mitochondrial membrane potential (ΔΨ_m_). Total membrane potential is significantly decreased in Pex5^cKO^ cells (H). Pex5^cKO^ cells show a more stable membrane potential under the stress protocol than WT cells (I, traces normalized at t=0). N = 40 cells from 3 mice per group. (E-I) Two-way ANOVA, Bonferroni’s multiple comparisons. (**J-M**) In isolated myocytes force development was determined by increasing the stimulation frequency from 1 to 4 Hz for 60 s and back after stepwise stretching the cell until the diastolic sarcomere length reached 2.0 µm. Example from WT (J). Pex5^cKO^ cells show less force development (K) compared to WT myocytes. Contraction velocity (L) and relaxation velocity (M) are significantly decreased in Pex5^cKO^ cells. WO = washout, BL = baseline. 6-9 cells from 3-4 mice per group, two-way ANOVA, Bonferroni’s multiple comparisons.

### Peroxisomes are essential for cardiac energetics and Ca^2+^ homeostasis

To understand the mechanistic details underlying the contractile dysfunction of the whole heart, we next monitored cellular energetic function. First, we analyzed the redox state of NAD(P)H and FAD in isolated cardiomyocytes in response to β-adrenergic and 0.5- to 5-Hz stimulation (Figure 7E-G and S7C). The level of the oxidized form of FAD was decreased in Pex5^cKO^ (Figure 7E,F). In consequence, the NAD(P)H/FAD ratio was more reduced at baseline in Pex5^cKO^ compared to WT (Figure 7G). During the workload transition via β-adrenergic and high frequency stimulation, ADP-induced acceleration of respiration induces oxidation of NAD(P)H and FADH_2_ (ref ^30,31^), like we observed in WT cardiomyocytes (Figure 7E-G and S7C). In PEX5^cKO^, however, the workload-induced increase in oxidized FAD was blunted (Figure 7E,F). In addition, NAD(P)H oxidation during workload transition was decreased in PEX5^cKO^, albeit not significantly (Figure S7C). The NAD(P)H/FAD ratio under workload conditions in Pex5^cKO^ recovered earlier compared to WT (Figure 7G and S7C). As a diminished oxidation of FAD might result from defective respiration, we assessed the mitochondrial membrane potential. When comparing resting myocytes, the mitochondrial membrane potential (ΔΨ_m_) was lower in Pex5^cKO^ vs. WT myocytes (Figure 7H), and its marginal dissipation (typically in response to ADP-induced acceleration of respiration) was less pronounced during the physiological stress protocol (Figure 7I).

The contractile dysfunction observed *in vivo* in the Pex5^cKO^ hearts together with the mitochondrial bioenergetic failure prompted us to take a deeper look into the contractile machinery on single cell level using force measurements in isolated cardiomyocytes. Preload-dependent force potentiation correlates to increased mitochondrial oxygen consumption (Frank Starling law)^32^. This energetic requirement is underestimated when measuring sarcomere shortening in myocytes without mechanical preload. We therefore stretched cardiomyocytes to physiological sarcomere lengths (∼2 µm) (Figure 7J and S7D,E). Under these conditions, diastolic sarcomere length was longer despite similar diastolic tension in Pex5^cKO^ vs. WT cardiomyocytes (Figure S7F,G). At a pacing frequency of 1 Hz we did not detect a difference in force between the groups, but when increasing pacing frequency to 4 Hz, Pex5^cKO^ cells develop substantially less systolic force (Figure 7K and S7H), implying peroxisome requirement upon increased energy demand. Also, we observed slower rates of contraction and relaxation indicating a reduced SERCA function similar to what we saw in the Pex5^iKO^ model (Figure 7L,MI). Hence, mechanical preload impairs systolic and diastolic function in Pex5^cKO^ mice and unmasks contractile deficiency on single cell level.

In line with this observation, we detected a trend towards axialization (loss of transversal elements, more axial orientation) of the T-tubule structures in our Cav3-stained isolated cardiomyocytes from the Pex5^cKO^ hearts (Figure 5F). These modifications in the T-tubule structure are often observed in HF and DCM and typically involve alterations in Ca^2+^ homeostasis and contractility ^33,34^.

### Peroxisomes compromised in a murine heart failure model

Our data show that absence of functional peroxisomes results in cardiac dysfunction, a finding that can be of importance for patients with milder forms of peroxisomal disorders or with any disease in which peroxisomes are indirectly affected^11,12^. We next wanted to ask, whether, vice versa, peroxisomes are affected in the failing heart. To address this question, we investigated peroxisomes in a mouse model of pressure overload-induced cardiac hypertrophy and HF.

We analyzed peroxisomes in mice with transverse aortic constriction (TAC) surgery and compared them with a sham control group. In TAC mice, pressure overload is induced by surgical constriction of the transverse aorta, limiting left ventricular outflow leading to cardiac hypertrophy and eventually HF^35^. One week after TAC surgery mouse hearts show clear signs of hypertrophy development with significant increase of left ventricular mass/body weight ratio (LVM/BW) (Figure 8A), increased septum thickness (Figure 8B) and increased relative wall thickness (Figure 8C). Further, we observed a significant increase in cardiomyocyte size one week after TAC operation indicating cellular hypertrophy (Figure 8E,F). As expected, atrial natriuretic peptide (ANP, *Nppa gene*) expression was upregulated (Figure S8A), indicative of increased hemodynamic load. However, the ejection fraction, the percentage of the total amount of blood that is pumped out with each heartbeat, was not significantly reduced (Figure 8D) meaning that cardiac function is still compensated. We stained peroxisomes in isolated cardiomyocytes from these mice, and Pex14 staining revealed a significant increase in peroxisome number (Figure 8G-I and S8B) compared to sham-operated hearts one week after surgery, while the peroxisome size was not affected (Figure 8J and S8B). Staining of the peroxisomal matrix protein Acaa1 in isolated cardiomyocytes after TAC surgery showed a similar phenotype (Figure S8C,D). Staining of peroxisomes in paraffin-embedded tissue samples from mouse hearts isolated after TAC confirmed significant increase in peroxisome number per area without change in size (Figure S8E,F). Further, we analyzed expression of peroxisome-related genes in hearts of sham-operated and TAC (one week) mice. Transcriptomic analysis (RNA seq) revealed significant mRNA downregulation of most peroxisomal proteins (Figure 8K and S8G-I). qPCR analyses of TAC- and sham-operated hearts confirmed significant mRNA downregulation of the peroxisomal proteins like the peroxisome proliferator-activated receptor alpha (*Ppara,* Figure 8L), the acyl-coA synthetase long chain family member 1 (*Acsl1*, Figure 8M), catalase (*Cat,* Figure 8N), hydroxysteroid dehydrogenase-like 2 (*Hsdl2,* Figure 8O), *Pex11a* (Figure 8P), *Gnpat* (Figure 8Q), and *Phyh* (Figure 8R). Interestingly, the levels of RNA encoding the peroxisomal membrane proteins *Abcd3*, *Pex14*, *Pex5*, Pex3, and *Pex16* were unchanged (Figure S8J). Only few genes were upregulated one week after TAC (Figure S8G) and many of them (*Nos2, Cyb5r3, Tmem173* and *Vim*) are already known to be associated with myocardial remodeling or HF. Hence, the elevated number of peroxisomes was not paralleled by elevated expression of peroxisomal proteins, suggesting compensatory upregulation of peroxisomal biogenesis after loss of matrix proteins. Eight weeks after TAC, hearts showed pronounced LV failure with significant increase in LVM/BW (Figure 8S), LV dilatation (Figure 8T) and decrease in ejection fraction (Figure 8U) and fractional shortening (Figure 8V; defined as the fraction by which the LV shortens during a cardiac cycle). The relative wall thickness and the cardiomyocyte size was unchanged between the groups (Figure S8K,L). Interestingly, peroxisome number and size in TAC cardiomyocytes (stained via Pex14 and Acaa1) were normalized and appeared unaltered when compared to cardiomyocytes from sham-operated animals (Figure 8W-Z and S8M). Also, downregulation of peroxisomal genes was less pronounced (Figure 8Zi and S8N-P). Hence, these findings show a dynamic response of peroxisomes to TAC-induced hypertrophy and HF.

**Figure 8:**
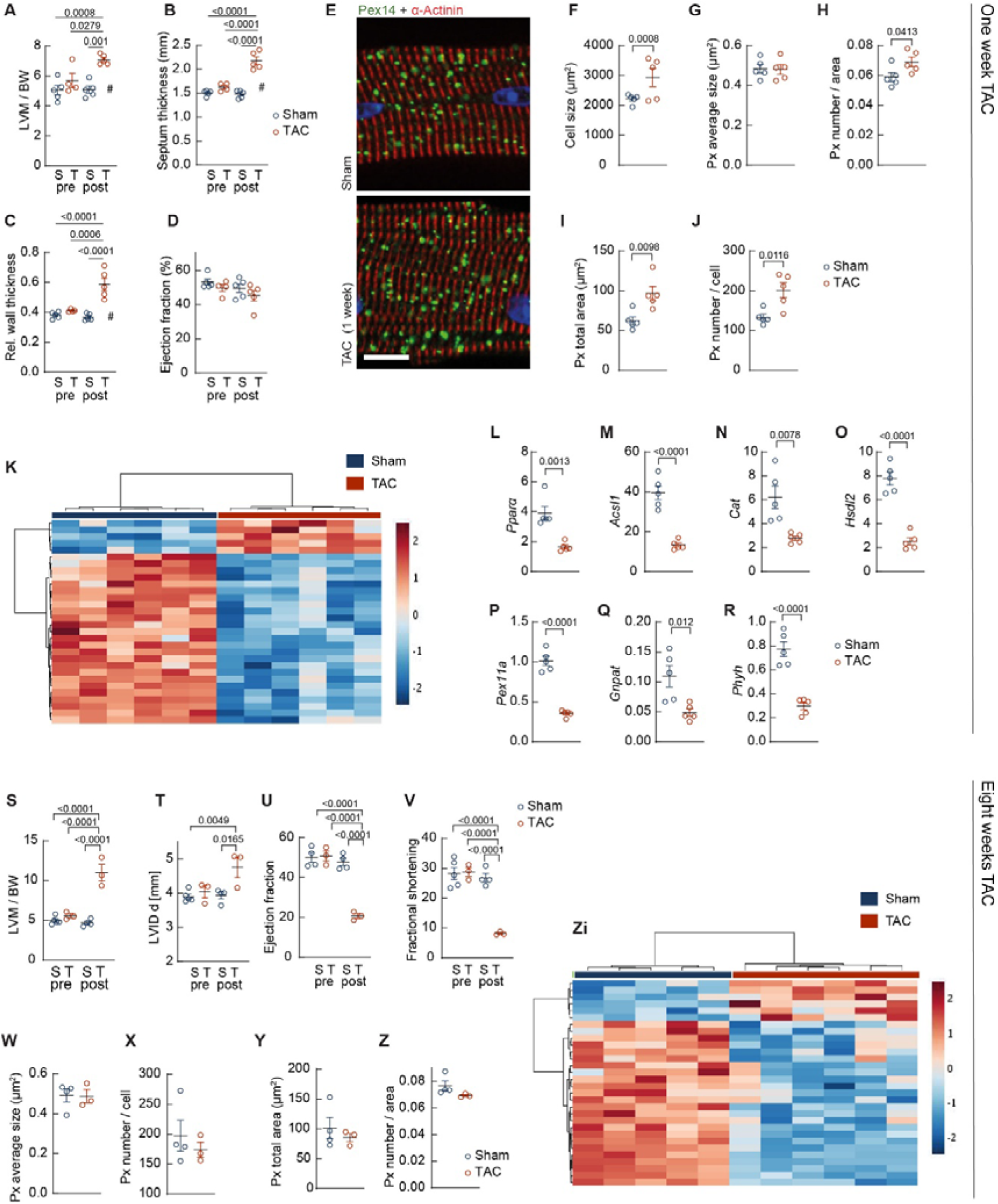
Peroxisome plasticity in the failing heart. (**A-J**) Peroxisome plasticity in cardiomyocytes from sham-operated mouse hearts and mouse hearts isolated one week after transverse aortic constriction (TAC). (A-D) Echo parameters from mice used in (E-J). N=5 mice per group, One-way ANOVA, Tukey’s multiple comparisons. LVM = left ventricular mass, BW = body weight. (E) Cardiomyocytes isolated from mouse hearts one week after TAC are hypertrophic and show peroxisome accumulation but no change in peroxisome size. Scale: 10 µm. (F-J) Quantification of (E). N = 5 mice per group. (**K**) Heatmap of mRNA Sequencing of mouse hearts one week after TAC operation shows significant downregulation of several peroxisomal genes (gene list derived from ^67^). (**L-R**) Verification of mRNA downregulation of selected genes from (K) and additional peroxisomal genes (one week after TAC, normalized to GUSB). N=5 mice. Px = peroxisome. (**S-V**) Echocardiographic analysis of WT mice before (pre) and eight weeks after TAC (post) shows significantly increased LVM / BW, dilation of the left ventricle and decreased ejection fraction and fractional shortening; N=4 sham and 3 TAC mice. One-way ANOVA. S = sham, T = TAC. (**W-Z**) Analysis of peroxisome staining of isolated CMs from sham and eight week-TAC mouse hearts. Peroxisomes were stained by anti-Pex14. No change in peroxisome size or peroxisome number was observed. N = 4 WT, 3 TAC mice. (**Zi**) Heatmap of RNA sequencing eight weeks after TAC.

In human and murine models, peroxisome dysfunction disrupts cardiac energy metabolism, mitochondrial structure, Ca^2+^ homeostasis and contractility. The resulting impaired force generation ultimately leads to HF. In the dilated mouse heart, compromised energy supply, especially in conditions of high demand, in combination with the defective Ca^2+^ homeostasis leads to irregular electrical conduction and thereby arrhythmic vulnerability, which may account for the apparently spontaneous death of Pex5cKO animals (Figure 9).

**Figure 9:**
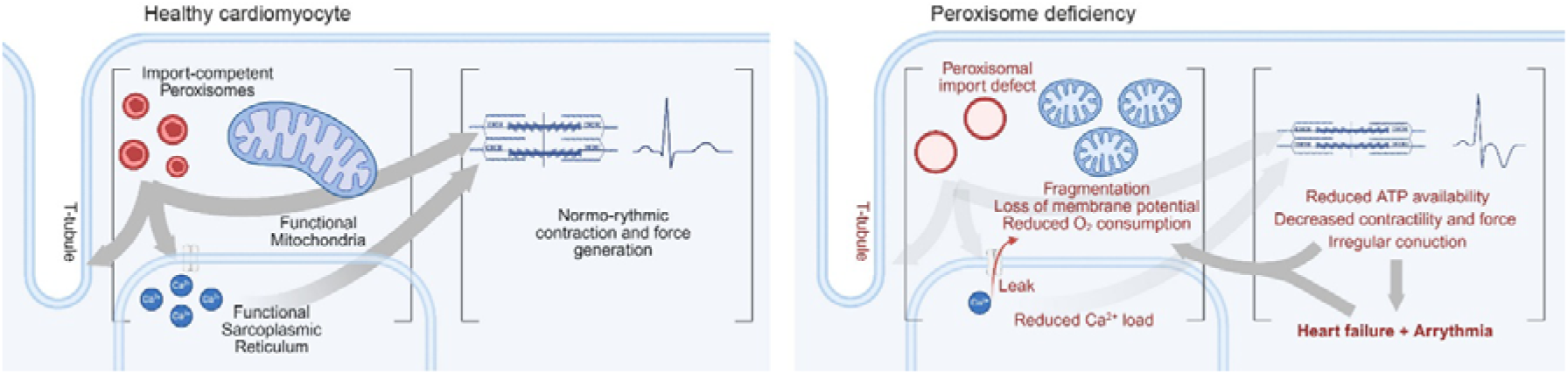
Peroxisome-dependent cardiac physiology and development of heart failure. Interorganellar crosstalk involving peroxisomes is a prerequisite for normal cardiac physiology in healthy cardiomyocytes. Loss of functional peroxisomes leads to mitochondrial aberrations with reduced respiration and disturbs SR Ca^2+^ homeostasis and T-tubule morphology. The disrupted organelle function results in irregular conduction, decreased contractility and force generation on cellular level, eventually leading to systolic and diastolic dysfunction, and finally failure of the whole organ. Vice versa, cardiac hypertrophy and heart failure trigger peroxisome responses. (Designed with BioRender.com).

## Discussion

The precise functions of peroxisomes in metabolism and protein homeostasis depend on the organism and the organ system. This becomes apparent when looking at patients and mouse models with peroxisomal biogenesis disorder which display a broad spectrum of dramatic consequences for several different organ systems^2^. Brain, liver, and kidney peroxisomes and the consequences of peroxisomal deficiency in these organs are well studied. Cerebro-hepato-renal symptoms are present from birth onwards and these patients usually die very young^36^. HF has been diagnosed in patients with milder PBD forms. However, the contribution of peroxisomes to heart physiology and pathology is still understudied, and we here dissect in detail the functional role of peroxisomes in cardiomyocyte function using engineered human heart tissue and two mouse models (Pex5^cKO^ and TAC).

Cardiac *PEX5* deficiency in human iPSC-CMs and the mouse heart leads to mitochondrial damage, disruption of mitochondrial membrane potential and deficits in oxygen consumption. This severe mitochondrial phenotype echoes a discovery made over 50 years ago while studying hepatocytes from Zellweger patients. In these cells, not only (functional) peroxisomes appeared to be absent, but mitochondria were also defective^11^. Since then, numerous studies have reported and analyzed peroxisome-induced mitochondrial dysfunction^37–41^. Mitochondrial damage is multifactorial, driven by metabolic factors such as the accumulation of cardiotoxic α-oxidation^26,42^ and β-oxidation substrates^41,43^ as well as protein mislocalization^38–40^ or loss of plasmalogens^44–46^. Recent work further shows a greater contribution of peroxisomal metabolism to cellular energy production than earlier models of peroxisome-mitochondria collaboration had suggested^12^.

The peroxisome-triggered respiratory deficit results in decreased total ATP production and this induces defects in Ca^2+^ homeostasis. Reduced post-rest potentiation and prolonged Ca^2+^ transient decay or slower rates of relaxation velocity, respectively, suggest reduced SERCA activity resulting in decreased SR Ca^2+^ load. Previous studies revealed that an energetic deficit primarily affects SERCA function and thereby, diastolic and systolic function and the contractile reserve in heart failure^47,48^. In fact, mechanical preloading revealed hypocontractility on a cellular level in the Pex5^cKO^ cardiomyocytes, which substantially increases the cells’ energetic requirements important for the Frank Starling mechanism^32^. These findings are to some extent reminiscent of our findings in a mouse model for Barth syndrome, a mitochondrial cardiomyopathy, in which sarcomere shortening was rather hyperdynamic in *unloaded* conditions, but hypocontractile when *mechanically preloaded* to physiological lengths, increasing the energetic demand which due to mitochondrial defects cannot be sufficiently matched^30^. Therefore, the mitochondrial dysfunction in Pex5^cKO^ myocytes and also in the human model likely contributes to contractile dysfunction.

ATP-deficiency and SR Ca^2+^ leak potentially additionally account for conduction defects in cardiac peroxisome-deficiency, as this can impair the function of cardiac ATPases inducing ion channel imbalances that affect the generation and propagation of electrical impulses in the heart^49^. Recently, we showed that peroxisomes are sites of intracellular Ca^2+^ handling in cardiomyocytes^50^. The extent and the mechanistic details of how peroxisomes contribute to cardiac Ca^2+^ homeostasis will be an exciting area for future studies.

Peroxisome-dependent plasmalogen deficiency is a contributor to conduction defects^51^. Peroxisomal ghosts in the *PEX5*-deficient cardiomyocyte can no longer import the two respective enzymes, dihydroxyacetone phosphate acyltransferase (DAP-AT, encoded by Gnpat) and alkyldihydroxyacetonephosphate synthase (AGPS), as seen in patients with PBD^52^ and in mouse models^53,54^. Plasmalogen biosynthesis appears to be vital for the heart^24^. Plasmalogens normally make up to 25% of the cardiac lipids and are essential for the structural integrity and fluidity of the myocytes^55^. Loss of plasmalogens leads to increased cellular susceptibility to ROS stress, which can contribute to HF development^56^. Plasmalogen loss and mitochondrial fragmentation have been associated with gap junction or conduction defects^51,57^, possibly explaining the observed pro-arrhythmic phenotype and finally the abrupt death of the Pex5^cKO^ mice. Conversely, peroxisomes contribute to ferroptosis via plasmalogen synthesis^58,59^. Future studies will show to what extent and by which molecular mechanism peroxisomal ether lipids contribute to the observed effects.

Dysregulation of peroxisomal factors after pressure overload implies a role of peroxisomes in cardiac physiology, especially under stress conditions. We further observed accumulation of peroxisomes shortly after TAC surgery, which was normalized eight weeks after TAC. These findings strengthen the model of a peroxisome-mitochondrial co-regulation, because in response to pressure overload, mitochondrial fragmentation and autophagy are transiently upregulated and normalized at later time points^60^. PEX5 has recently been described as a protector of cardiac hypertrophy in rat cardiomyocytes^61,62^. Our findings suggest that peroxisome activation plays a role in left ventricular hypertrophy and consequent HF development. The Pex5^cKO^ mice appear healthy at birth and DCM development starts not before five weeks of age. The Pex5 HF phenotype develops slowly, continuously and initially unnoticed, similar to human compensated HF^63^. In contrast to that, the neurological phenotypes present in Zellweger syndrome are present from birth on and lead to death of patients in early childhood^36^. Due to this serious neurological phenotype and the early death of PBD patients, the significant contribution of peroxisomes to cardiac function, manifesting as a later-onset cardiomyopathy (beyond the first years of life), is usually masked. Conversely, patients suffering from Refsum disease, a peroxisomal (biogenesis) disorder that can be caused by single enzyme defect or deficiency in the PEX5-coreceptor, are typically diagnosed with cardiomyopathy; and HF is the most frequent cause of death in this patient cohort^6^.

The models established in this work can be used to dissect the relative involvement of peroxisomal functions, protein homeostasis, and organelle interaction to cardiac health and HF development. In conclusion, our findings highlight the essential and previously overlooked role of peroxisomes in the heart, potentially serving as a new target for intervention.

## Materials and Methods

### Animals

Floxed *Pex5* mice (*Pex5*^fl/fl^) were kindly provided by Myriam Baes and Myh6-Cre mice were kindly provided by Karl Toischer. Myh6-Cre^+/-^ mice were crossed to BL/6N-*Pex5*^fl/fl^ mice to specifically knockout *Pex5* from cardiomyocytes (Pex5^cKO^ = Myh6-Cre^+/-^-Pex5^fl/fl^). Myh6-Cre^-/-^-Pex5^fl/fl^ (=WT) and Myh6-Cre^+/-^-Pex5^+/+^ (=Cre-control, WT^c^) were used as control groups. Genotypes were determined via polymerase chain reaction analysis on ear DNA using the Myh6-Cre primer sets (for: 5′-ATG ACA GAC AGA TCC CTC CTA TCT CC-3′, rev: 5′-CTC ATC ACT CGT TGC ATC ATC GAC-3′ and for: 5′-CAA ATG TTG CTT GTC TGG TG -3′, rev: 5′-GTC AGT CGA GTG CAC AGT TT -3′) and Pex5 primers (for: 5′-TCT GGT TCC CAT TGG CCA GGG TGG C-3’, rev: 5′-CGG GGA GTA CGA CAA GGC TGT GGA CT-3′). Heart and body weight were determined in mice used for echocardiography measurements. C57Bl/6N (Charles River Laboratories) mice were used for transaortic constriction (TAC) operations. Mice were housed in a 12 h light/dark cycle at 21°C and fed ad libitum on standard chow rodent diet (Ssniff Spezialdiaeten GmbH). Animal experiments were approved by the animal ethics committee (AZ 17/2412, RUF 55.2-2-2532-2-659) and conducted in accordance with institutional guidelines.

### Transaortic constriction and echocardiography

TAC was done as previously described^64^. Briefly, the intervention was performed after anaesthesia by tying a braided 5-0 polyviolene suture ligature around the aorta and a blunted 27-gauge needle and subsequent removal of the needle. Sham controls underwent the same procedure but the suture was not tied. For echocardiography, mice were anesthetized by 2.4 % isoflurane inhalation and ventricular measurements were done with a VisualSonics Vevo 2100 Imaging System equipped with a MS400 30 MHz MicroScan transducer as described in Ref^64^. All TAC and echocardiography procedures were performed by the SFB 1002 S01 service unit.

### Histology

Mice were anesthetized by isoflurane and sacrificed by cervical dislocation. Mouse hearts were removed, washed in physiological saline and fixed in 10 % phosphate-buffered formaldehyde for 48 h. Hearts were washed in H_2_O overnight, dehydrated and embedded in paraffin. 3 µm slices were mounted on microscope slides and stained with HE or Picrosirius or with the respective antibodies (listed in Table S1) and antigen retrieval (Fa Dako D11828 pH 9.0 1:50). Primary antibodies were either visualized by secondary fluorescent antibodies (listed in Table S1) or HRP-coupled secondary antibodies and DAB substrate (Dako K3468).

Isolation and immunofluorescence staining of mouse ventricular cardiomyocytes Isolation of ventricular myocytes for immunofluorescence imaging was performed as described in ref^33^. Ventricular cardiomyocytes from mice hearts were isolated by retrograde perfusion of the heart according to the Langendorff technique after isoflurane anaesthesia cervical dislocation. After that, ascending aorta of hearts was cannulated and mounted in a Langendorff perfusion apparatus. Hearts were perfused with 4 mL/min at 37°C. After initial perfusion with Ca^2+^-free Tyrode’s solution for 2 min, hearts were digested with collagenase type II (610 U/ml) and 40 µM CaCl_2_ in Tyrode’s solution for 7 min at 37°C. Following perfusion, the hearts were minced in the digestion buffer. The tissue was carefully resuspended with a 10 mL serological pipette and the digestion process was stopped by adding stop buffer before resuspending again. The cardiomyocytes were washed three times with stop buffer. Thereafter, cells were allowed to sediment by gravity for eight min at room temperature.

For immunofluorescence staining of cardiomyocytes, cells were seeded on laminin-coated cover slips and incubated for 1 h at room temperature or 30 min at 37°C. Fixation was performed with 4 % paraformaldehyde for 10 min followed by 2 x 5 min washing with PBS. Cells were incubated with blocking solution (5 % BSA, 0.5 % Triton X-100 in PBS) for 1 h. Cells were then incubated with the first antibody (listed in Table S1) diluted in blocking solution at 4°C overnight. Cells were washed with PBS 3 x 10 min, incubated with the second antibody (listed in Table S1) diluted in blocking solution for 3 h at room temperature and mounted with Vectashield mounting medium.

### Microscopes and image analysis

Cells were imaged by widefield microscopy (Axio Observer Z1 equipped with Zeiss Colibri 7 and with 63× oil Fluar; deconvoluted). When not stated otherwise, widefield microscopy and deconvolution was used for imaging. Linear contrast enhancements were done with ImageJ. Analysis and image processing was done using Fiji (http://fiji.sc/) according to ImageJ User Guide. To count peroxisomes in images, auto threshold of MaxEntropy was applied on cell overview images of the respective staining. Peroxisomes were then counted automatically using the built-in function Analyse Particles. Particles with size smaller than 0.03 µm² were excluded from analysis. Peroxisome morphology parameters area, roundness and solidity were analyzed doing the same steps but on the five-times zoomed in confocal and STED images.

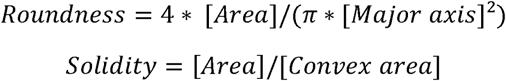

3D structured illumination microscopy was performed on an OMX v4 microscope (GE Healthcare). Immediately before imaging, cells were incubated with MitoTracker Green (Invitrogen) for 30 min, and Hoechst 33342 (Invitrogen) for 5 min at 37°C. Images were reconstructed and registered with softwoRx 7.0 (GE Healthcare). Quantitative analysis of mitochondrial morphology was performed with Mitochondria analyser for ImageJ^65^.

### Protein expression, plasmalogen measurement, mRNA expression and sequencing

Mice were anesthetized by isoflurane and sacrificed by cervical dislocation. For protein expression analysis, the ventricular part of the heart was separated from the atrial part, transferred into reaction tubes and frozen in liquid nitrogen. After thawing on ice, the cardiac tissue was homogenized 5 times for 5 sec. The homogenate was centrifuged two times at 1,300 x g for 10 min at 4 °C. Supernatants were transferred into a new reaction tube. Proteins were detected in homogenates of the ventricular heart tissue by SDS-PAGE and Western blot (antibodies listed in Table S1). Anti-Gapdh was used as loading control. All uncropped WB images are shown in S9.

For plasmalogen measurements, heart tissue was homogenized in PBS with 2 mM EDTA (5 mL/g), centrifuged 10 min at 14000 rpm at 4°C. Supernatant and pellet were stored at - 80°C. The Pellet was dissolved in 500 µl buffer per 50 mg tissue. 50 µg was used for plasmalogen measurement.

For RNA isolation, mouse ventricular tissue was macrodissected. RNA was isolated using peqGOLD TriFast (Quiagen). Nucleic acid was quantified by a Nanodrop photometer (Thermo Scientific). 2 µg RNA was used for cDNA synthesis, together with 1 µg Oligo(dT)20 primer and 100 U M-MLV reverse transcriptase (Promega) for 1 h at 42°C. Quantitative real-time PCR (qPCR) analyses were performed with SYBR Green (Promega) on a QuantStudio 3 real-time cycler (ThermoFisher) using primers listed in Table S2. Gene expression was normalized to GAPDH or GUSB (glucuronidase beta).

RNA Sequencing data from FVBN mice were taken from Ref^66^. Peroxisomal gene list was derived from Ref^67^. Genes with a score below 5 were omitted from the list. Additionally, Acbd3, Pomc, Sel1L, Idi2, Drp1, Fis1, Mff, Gdap, Miro1, Pparα, PparL and Pparδ were added to the list. The module “Statistical Analysis” of the online freeware Metaboanalyst (version 4.0) was used for analysis.

### Mitochondrial fractionation and Blue Native PAGE

For mitochondrial isolation, tissue was minced in isolation buffer (20 mM HEPES pH7.6, 220 mM Mannitol, 70 mM Sucrose, 1 mM EDTA) with freshly added 0.5 mM PMSF, complete protease inhibitors (Roche) and phosphatase inhibitors (Roche) at 4°C, and tissue was homogenized using a manual potter. Tissue debris was removed by centrifugation for 15 min at 800 g. The pellet was homogenized again, and the supernatant was transferred to a new tube further cleared by repeated centrifugation steps. Collected supernatant was centrifuged at 10000 g for 10 min to harvest mitochondria. The mitochondria pellet was washed and centrifuged again at 10000 g for 10 min. Mitochondrial total protein content was determined using the DC protein assay from BioRad according to manufacturer’s instructions. For BN-PAGE analysis, mitochondria were solubilized in digitonin buffer (20 mM Tris-HCl 20, 50 mM NaCl, 0.1 mM EDTA, 2 mM PMSF, 10 % glycerol (v/v) 1 % digitonin (w/v); pH 7.4) for 15 minutes on ice. After a clarifying spin, the samples were separated on a 4-13 % BN-PAGE. Native protein complexes were transferred on a PVDF membrane followed by immunoblotting with specific antibodies.

### Unloaded membrane-intact cardiomyocytes

For cardiomyocyte isolation, mice received buprenorphin (0.1 mg/kg) and heparin (500 IU/kg) and were anesthetized with isoflurane (5% at 0.5 L/min). The heart was excised and processed when mice became insensible to pedal reflex. Cardiomyocytes were isolated from mouse hearts by enzymatic digestion as previously described^68,69^. In brief, experiments with intact cardiac myocytes were performed at 37°C. During the experiment, cells were continuously perfused with Normal Tyrodés (NT) solution containing 130 mM NaCl, 5 mM KCl, 1 mM MgCl_2_, 1 mM CaCl_2_, 10 mM Na-HEPES, 10 mM glucose, 2 mM sodium pyruvate, 0.3 mM ascorbic acid and pH-control at pH 7.4. Unloaded membrane-intact cardiomyocytes were field-stimulated at 0.5 Hz for 2 minutes. A simulated increase in cardiac workload (i.e., stress protocol) was conducted, where cells were subsequently perfused with NT solution containing the pharmacological β-adrenergic stimulator isoprenaline (Iso, 30 nM), and when the Iso-induced increase in contractility was detected, stimulation frequency was increased to 5 Hz for 3 minutes. Stimulation at 0.5 Hz was resumed with NT (wash-out (WO) period) until contractility was again comparable to baseline conditions. During the experiments, NAD(P)H and FAD autofluorescence was recorded. Calibration of NAD(P)H and FAD signals was conducted with 5 µM carbonyl cyanide-p-trifluoromethoxyphenylhydrazone (FCCP) and cyanide (4 mM) to induce the complete oxidation and reduction, respectively, of the mitochondrial pyridine nucleotide pools. ΔΨ_m_was measured by incubating cells with 1µM TMRM for 10 minutes at 25°C (excitation 540 nm; emission 605 nm).

### Loaded membrane-intact cardiomyocytes

To monitor force development upon variations in length in isolated cardiac myocytes (i.e., cardiomyocyte work output), we used the established IonOptix force transducer as previously described^68,70^. Cardiomyocytes were perfused with NT solution containing 2 mM CaCl_2_. In brief, both the cantilever of the force transducer and the piezo motor are initially embedded with “precoat” for a few seconds and thereafter coated with “MyoTac”. The holders were then transferred into the chamber, in which the cardiomyocytes were placed on a 2 % BSA-laminated cover slip. Normal beating cells in the correct plane orientation of both cantilevers were attached and then stretched by the piezo motor up to a sarcomere length of 1.9 µM. Measurements were performed at room temperature. Stimulation frequency was 1 Hz and 4 Hz.

### Cardiac electrograms

Hearts from WT and Pex5^cKO^mice were quickly extracted after cervical dislocation and immediately placed in ice-cold Dulbecco’s phosphate-buffered solution. Non-cardiac tissue was discarded, and the heart perfused via the aorta with Tyrode’s solution (1.8 mM CaCl_2_, 140 mM NaCl, 5.4 mM KCl, 2 mM MgCl_2_, 10 mM Glucose, 10 mM HEPES; pH adjusted to 7.4 using NaOH, bubbled with 100 % O_2_ at 37°C). A bipolar epicardial electrogram was obtained using a silver electrode in contact with the right atrium and a metal spoon at the heart’s apex. The signal was amplified with an Animal Bio Amp (FE136), recorded with PowerLab 16/35 and analyzed with LabChart 8.1.18 software (all ADinstruments). Baseline heart rate, QRS, and QT intervals were quantified during stable sinus rhythm (5-10 mins after starting perfusion) by averaging 25 heart beats with LabCharts ECG Analysis module (v2.4). Electrical stimulation (2 mA, biphasic, 2 ms) was triggered by the Powerlab 16/35 and applied with the Isolated Pulse Stimulator 2100 (A/M Systems, Washington, US) via bipolar silver electrodes at the basis of the right ventricle. The local effective refractory period (ERP) was analyzed as described previously ^71^. Briefly, we used an S1-S2 protocol, with 6 stimuli at a cycle length of 150 ms (S1) followed by a premature stimulus (S2) with a variable delay (<150 ms) that was decreased in 5 ms steps. The shortest time period with successful S2 pacing was defined as the local ERP.

### Generation and cardiac differentiation of human iPSC lines, calcium and TMRE measurement

Human iPSC lines from a healthy donor and a CRISPR/Cas9-engineered *PEX5*-deficient line were used in this study. WT hiPSC line UMGi014-C clone 14 (isWT1.14) was generated from dermal fibroblasts using the integration-free Sendai virus as described previously^72^. Homozygous insertion of the pathological gene variant PEX5 c.397C>T/p.Q133X in the WT hiPSC line UMGi014-C clone 14 was performed using ribonucleoprotein (RNP)-based CRISPR/Cas9 by targeting exon 5 of the PEX5 gene, as previously described^73^. The guide RNA target sequence was (PAM in bold): 5’-AGATG CTGTG GATGT AACTC **AGG**-3’. For homology-directed repair, a single-stranded oligonucleotide with 45-bp homology arms was used. After clonal singularization, successful genome editing was identified by Sanger sequencing and the CRISPR-modified isogenic hiPSC line UMGi014-C-7 clone 39 (isWT1-PEX5-KO.39) underwent pluripotency characterization as well as chromosomal integrity by digital karyotyping, as previously described^74^. Human iPSC lines were differentiated into ventricular cardiomyocytes via WNT signaling modulation and subsequent metabolic selection and cultured for at least 30 days, as previously described^75^. Human iPSCs and derived cardiomyocytes were cultured in feeder-free and serum-free culture conditions in a humidified incubator at 37 °C and 5 % CO_2_. The study was approved by the Ethics Committee of the University Medical Center Göttingen (approval number: 10/9/15) and carried out in accordance with the approved guidelines. Following differentiation, purity of hiPSC-CMs was determined by flow cytometry analysis using an antibody against cTNT (Suppl. Table S1). Cytosolic Ca^2+^ recordings in hiPSC-CMs were performed as previously described^74^. For measurement of mitochondrial membrane potential, cells were incubated with TMRE for 30 min, washed with prewarmed PBS and fluorescence intensity was measured in a Tecan infinite 200Pro Plate reader at 549/575 nm. TMRE signal was normalized to MitoTracker Green (200 nM, measured at 490/516 nm).

### Cellular respiration analysis

Mitochondrial respiration and changes in cellular metabolism were characterized by extracellular flux analysis using an Agilent Seahorse XFe24 Analyzer (Agilent Technologies). The mitochondrial stress test was performed on 8-week-old iPSC derived cardiomyocytes (WT and *PEX5*-deficient) by measuring the oxygen consumption rate (OCR) in presence of electron transport chain inhibitors. Briefly, the iPSC-CMs were seeded one week prior to the assay and cultured in Seahorse XFe24 Cell Culture microplates. One day before the assay, the Seahorse XF Sensor Cartridge was hydrated by adding 1 mL of the Seahorse XF Calibrant Solution to each well and then kept in a non-CO_2_ incubator at 37 °C for at least 24 h. On the day of the assay, the cells were washed twice with Agilent XF RPMI without any additives, then kept in 1 mL XF Assay Medium (Agilent XF RPMI supplemented with 10 mM glucose, 1 mM pyruvate, 2 mM glutamine, 200 µM L-Asc and 2 % B27 supplement) per well for 1 h at 37 °C in a non-CO_2_ incubator. The oxygen consumption rate (OCR) was assessed following sequential injections of ETC inhibitors diluted in XF Assay Medium (with final concentration in wells): oligomycin (2 µM), FCCP (2 µM), Rotenone and Antimycin A (both at 1 µM). Subsequent cell quantification for normalization was performed via nuclei counting using an automated cell imager (CELLAVISTA, Synentec, Germany) after staining the cells with Hoechst 33342 (10 µg/mL).

### Immunofluorescence staining and protein expression in hiPSC-CMs

For analysis of protein expression, hiPSC-CMs were washed with PBS and homogenized on ice in RIPA buffer. After centrifugation (5 min at 14000 rpm at 4°C), protein concentration was determined by BCA assay. Homogenates of hiPSC-CMs were then used for SDS-PAGE and Western blot (antibodies listed in Suppl. Table S1). Anti-GAPDH antibody was used as loading control. All uncropped WB images are shown in S6. For immunofluorescence staining, hiPSC-CMs were washed with PBS, fixed for 20 min in 4% paraformaldehyde, permeabilized with 0.5 % Triton X100 in PBS for 5 min and blocked in 10 % BSA in PBS for 1 h. Antibodies were diluted in blocking buffer. Primary antibody staining was carried out overnight at 4°C, secondary antibody for 1 h at room temperature and mounting was done with ProlongGOLD with Dapi.

### Generation and analysis of EHM

Human iPSC lines UMGi014-C clone 14 and isogenic UMGi014-C-7 clone 39 were used for EHM experiments. Defined, serum-free EHM was generated from hiPSC-CMs at day 30 and human foreskin fibroblasts (ATCC) at a 70:30 ratio according to previously published protocols^15,76^. Optical analysis of spontaneously beating EHM on flexible posts in a 48-well plate (myrPlate-TM5, myriamed GmbH) was performed on day 14, day 29, and day 43 of culture using a custom-built setup with a high-speed camera (Basler acA3088; 50 frames per seconds). Fully parallel online pole tracking was achieved by automated thresholding and center of gravity analysis of each well using custom C++ Software. Resulting traces were analyzed with a custom Matlab software for peak detection and analysis. Auxotonic contractions were quantified as EHM shortening (%) = (baseline-peak)/baseline pole distance. Isometric force measurements on day 43 of EHM culture were performed in thermostat-equipped organ baths (Föhr Medical Instruments) in Tyrode’s solution (120 mM NaCl, 1 mM MgCl2, 0.2-4 mM CaCl2, 5.4 mM KCl, 22.6 mM NaHCO3, 4.2 mM NaH2PO4, and 5.6 mM glucose; pH 7.4 was maintained by gassing with 5 % CO_2_/95 % O_2_) as previously described ^76^. EHM were electrically stimulated at 1.5 Hz with 5-ms square pulses of 150 mA. EHM were mechanically stretched at intervals of 125 μm until the maximum systolic force amplitude was observed according to the Frank-Starling law. Force-frequency response and post-rest potentiation were assessed at individual EC_50_ Ca^2+^ concentrations. For post rest potentiation, EHM electrical stimulation was resumed with 1 Hz after a 10 sec pause following 3 Hz stimulation. The first contraction after the pause was compared to baseline 1 Hz contractions ∼90 sec after restarting 1 Hz stimulation. hiPSC-CMs before EHM generation or after enzymatic dispersion of EHM were fixed in ice-cold 70 % ethanol and stained with antibodies (Suppl. Table 1) as previously described ^76^.

### Optical Mapping Experiments

To study action potential duration and conduction properties EHMs were incubated with the voltage-sensitive dye Di-4-Anepps (D1199, Invitrogen) for 10 minutes at 37°C^77^. Imaging was performed using a complementary Metal-Oxide semiconductor (CMOs) camera (Micam Ultima 10 x 10 mm² sensor, Sci-Media). During experiments EHM were maintained at 37 °C in carbonated bath solution (5 % CO_2_ and 95 % O_2_) containing: NaCl 126.7 mM, KCl 5.4 mM, MgCl_2_ 1.1 mM, CaCl_2_ 1.8 mM, NaHPO_4_ 0.42 mM, NaHCO_3_ 22 mM, glucose 5.5 mM, pH=7.45. EHM were electrically point-stimulated using a custom-built platinum/iridium bipolar electrode at stimulation frequencies between 0.5 and 3 Hz. The voltage indicator was excited with peak wavelengths of 531 nm using an LED (LEX3 illumination system, Sci-Media), and emission was passed through a 600 nm long-pass filter. A custom designed software (kindly provided by Prof. Dr. Igor Efimov, George Washington University) was used for analysis, generation of activation maps, and conduction velocity measurements. All optical mapping experiments were performed after initial incubation with blebbistatin for 15 min (5 µM, Sigma-Aldrich) to prevent motion artifacts.

### Statistical analyses

Statistical analysis was done with Excel or GraphPad Prism 8 by Student’s t-test or two-way ANOVA for repeated measurements. When multiple cells per mouse were measured nested ANOVA or nested t-tests were performed. P-values less than 0.05 were considered statistically significant. When not stated otherwise, two-sided, unpaired Student’s t-test was used. Further details are found in the accompanying figure legend.

## Acknowledgments

The authors thank Miriam Baes for Pex5^fl/fl^ mice. We thank Laura Cyganek, Nadine Gotzmann, Yvonne Hintz, Lisa Schreiber, and Yvonne Wedekind for technical assistance in hiPSC generation, genome editing and cardiac differentiation and Regina Waldmann-Beushausen for technical assistance with histology. We thank Marcel Zoremba and Robert Blume for echocardiography and Beate Knocke for echocardiography analysis (all SFB 1002 S01). We thank Irmgard Cierny, Anna Grönke, Annette Berbner, Manuela Erk and Johanna Heine for general technical assistance, Ralph Krätzner and Monika Schneider for plasmalogen measurements, Ignacio Lobos for help with image analysis, Eva Wagner and Gregory Antonius for help in STED imaging, Eric Buchholz for NGS data and discussions, Eric Schoger and help with Langendorff perfusion and Gabriela Salinas-Riester, Moritz Schnelle and Vivek Venkataramani for discussions.

## Author contributions

JH: Imaging and quantification of EHM sections, iPSC-CM characterization by IF and WB and quantification, Supervision of TAC experiments and analysis of TAC phenotype, CM isolation, staining, quantification, WB of heart lysates, evaluation and interpretation of data, drafting of manuscript and figure design; MT: EHM generation and characterization, figure, manuscript review, funding; YH: STED imaging and quantification; KB: Establishment of mouse breeding, Pex5 PCR for KO verification, imaging; AW: Electrophysiological characterization of cardiac electrograms (mouse hearts); OGG: iPSC-CM differentiation and Ca^2+^ measurements; RS: Optical mapping experiments; LT: Analysis of mitochondrial phenotype in iPSC-CMs; NL: Data interpretation; MW: iPSC-CM differentiation and analysis by WB and IF; PT: Mouse experimentation; MG: WB of mitochondrial proteins; WH: SIM imaging; TH: SIM imaging; BB: EHM generation; MC: EHM generation and characterization; AU: EM imaging of mouse hearts; SaS: Interpretation of mouse data, funding; KSB: manuscript review, funding; ME: Support with TAC experiments, analysis and interpretation of echocardiography data, interpretation of results; KT: Support with TAC experiments, interpretation of results; MK: Force measurement in mouse CM, autofluorescence of NAD(P)H and FAD, mitochondrial membrane potential, manuscript review; JD: Blue native and WB of mitochondrial complexes and proteins, funding; WHZ: Supervision of EHM and echocardiography experiments, interpretation of data, draft revision, funding; NV: Supervision and interpretation of optical mapping experiments, manuscript review, funding; TB: Electrophysiological characterization of cardiac electrograms (mouse hearts), manuscript review, funding; CM: Interpretation of force measurements, manuscript review; LC: Supervision of CRISPR editing and iPSC generation, analysis and interpretation and Ca^2+^ measurements in iPSC-CMs, figure, manuscript review, funding; ST: Study design and coordination, supervision, data analysis and interpretation of results, manuscript writing, funding

## Sources of Funding

This work was supported by the following grants: Deutsche Forschungsgemeinschaft (DFG) Collaborate Research Council ‘Modulatory units in heart failure’ SFB 1002/3 TP A10 to ST, TP C04 to MT and WHZ, TP A13 to NV, TP A14 to TB, TP D04 to KT and TP S01 to WHZ and LC; DFG TH 1538/3-1 to ST; DFG VO 1568/3-1 and VO1568/4-1 to NV, DFG-471241922 to KSB and SO 1223/4-1 to SaS; TO 822/1-1 to KT; the Fondation Leducq 20CVD04 to WHZ; TO 822/1-1 to KT; the Eva-Luise and Horst Köhler Foundation to ST; German Centre for Cardiovascular Research (DZHK) 81X4300105 and 81X2300198 to ST, DZHK 81X4300102, 81X4300115, 81X4300112 to NV, DZHK 81Z0300116 to WHZ and LC, DZHK 81Z0300119 to WHZ and DZHK doctoral stipend to AW; and the German-Israeli Foundation for Scientific Research and Development (GIF, grant number 1459) to ST. The F. Thyssen Foundation (Az 10.19.2.026MN) and DFG (471241922 and CRC1213) to KSB and SaS. DFG Ma 2528/8-1 (project 50580539), Ma 2528/9-2 (project 315254101) and SFB 1525 (project 453989101) to CM. DAAD PhD stipend ID 91572398 to YH. Marlies und Herbert Repkow Stiftung to ST. NV and WHZ were funded by Germany’s Excellence Strategy: EXC 2067/1—390729940. NL was funded by the German Research Foundation Excellence Strategy EXC-2049-390688087. JD was funded by the DFG (SFB1525, TP B5) and the Interdisziplinäres Zentrum für Klinische Forschung (IZKF; E-457). TB was funded by Germany’s Excellence Strategy: EXC 2067/1 (390729940).

## Competing interests

WHZ is founder and equity holder of Myriamed GmbH with an interest in drug development in human iPSC-based cell and tissue models. MT is advisor of Myriamed GmbH. NL is a member of the scientific advisory board of Trace Neuroscience.

## Data availability

All data supporting the findings of this study are available within the paper and its Supplementary Information.

## References

1. De Duve, C. & Baudhuin, P. Peroxisomes (microbodies and related particles). Physiological Reviews 46, 323–357 (1966).

2. Wanders, R. J. A., Baes, M., Ribeiro, D., Ferdinandusse, S. & Waterham, H. R. The physiological functions of human peroxisomes. Physiol Rev 103, 957–1024 (2023).

3. Kumar, R., Islinger, M., Worthy, H., Carmichael, R. & Schrader, M. The peroxisome: an update on mysteries 3.0. Histochem Cell Biol 161, 99–132 (2024).

4. Gärtner, J., Rosewich, H. & Thoms, S. The Peroxisome Biogenesis Disorders. in The Online Metabolic and Molecular Bases of Inherited Disease (eds. Valle, D. L., Antonarakis, S., Ballabio, A., Beaudet, A. L. & Mitchell, G. A.) (McGraw-Hill Education, New York, NY, 2019).

5. Thoms, S. & Erdmann, R. Peroxisomal matrix protein receptor ubiquitination and recycling. Biochim Biophys Acta 1763, 1620–1628 (2006).

6. Wanders, R. J. A. & Komen, J. C. Peroxisomes, Refsum’s disease and the α- and ω-oxidation of phytanic acid. Biochemical Society Transactions 35, 865–869 (2007).

7. Lopaschuk, G. D., Karwi, Q. G., Tian, R., Wende, A. R. & Abel, E. D. Cardiac Energy Metabolism in Heart Failure. Circ Res 128, 1487–1513 (2021).

8. Aksentijevic, D. et al. Mechano-energetic uncoupling in heart failure. Nat Rev Cardiol 22, 773–797 (2025).

9. Soliman, K., Göttfert, F., Rosewich, H., Thoms, S. & Gärtner, J. Super-resolution imaging reveals the sub-diffraction phenotype of Zellweger Syndrome ghosts and wild-type peroxisomes. Sci Rep 8, 7809 (2018).

10. Santos, M., Imanaka, T., Shio, H., Small, G. & Lazarow, P. Peroxisomal membrane ghosts in Zellweger syndrome--aberrant organelle assembly. Science 239, 1536–1538 (1988).

11. Goldfischer, S. et al. Peroxisomal and mitochondrial defects in the cerebro-hepato-renal syndrome. Science 182, 62–64 (1973).

12. Violante, S. et al. Peroxisomes can oxidize medium- and long-chain fatty acids through a pathway involving ABCD3 and HSD17B4. FASEB J 33, 4355–4364 (2019).

13. Tiburcy, M. & Zimmermann, W.-H. Modeling myocardial growth and hypertrophy in engineered heart muscle. Trends Cardiovasc Med 24, 7–13 (2014).

14. Jebran, A.-F. et al. Engineered heart muscle allografts for heart repair in primates and humans. Nature 639, 503–511 (2025).

15. Tiburcy, M., Meyer, T., Liaw, N. Y. & Zimmermann, W.-H. Generation of Engineered Human Myocardium in a Multi-well Format. STAR Protoc 1, 100032 (2020).

16. Holubarsch, C. et al. Existence of the Frank-Starling mechanism in the failing human heart. Investigations on the organ, tissue, and sarcomere levels. Circulation 94, 683–689 (1996).

17. Puglisi, J. L., Negroni, J. A., Chen-Izu, Y. & Bers, D. M. The force-frequency relationship: insights from mathematical modeling. Adv Physiol Educ 37, 28–34 (2013).

18. Hasenfuss, G. et al. Relation between myocardial function and expression of sarcoplasmic reticulum Ca(2+)-ATPase in failing and nonfailing human myocardium. Circulation Research 75, 434–442 (1994).

19. Mulieri, L. A., Hasenfuss, G., Leavitt, B., Allen, P. D. & Alpert, N. R. Altered myocardial force-frequency relation in human heart failure. Circulation 85, 1743–1750 (1992).

20. Agah, R. et al. Gene recombination in postmitotic cells. Targeted expression of Cre recombinase provokes cardiac-restricted, site-specific rearrangement in adult ventricular muscle in vivo. J Clin Invest 100, 169–179 (1997).

21. Baes, M., Dewerchin, M., Janssen, A., Collen, D. & Carmeliet, P. Generation of Pex5-loxP mice allowing the conditional elimination of peroxisomes. Genesis 32, 177–178 (2002).

22. Baes, M. et al. A mouse model for Zellweger syndrome. Nat Genet 17, 49–57 (1997).

23. Kunze, M. et al. Mechanistic insights into PTS2-mediated peroxisomal protein import: the co-receptor PEX5L drastically increases the interaction strength between the cargo protein and the receptor PEX7. J Biol Chem 290, 4928–4940 (2015).

24. Plessner, M., Thiele, L., Hofhuis, J. & Thoms, S. Tissue-specific roles of peroxisomes revealed by expression meta-analysis. Biol Direct 19, 14 (2024).

25. Nagan, N. & Zoeller, R. A. Plasmalogens: biosynthesis and functions. Progress in Lipid Research 40, 199–229 (2001).

26. Grings, M. et al. Phytanic acid disturbs mitochondrial homeostasis in heart of young rats: a possible pathomechanism of cardiomyopathy in Refsum disease. Mol Cell Biochem 366, 335–343 (2012).

27. Park, H. et al. Peroxisome-derived lipids regulate adipose thermogenesis by mediating cold-induced mitochondrial fission. J Clin Invest 129, 694–711 (2019).

28. Rehmani, T., Salih, M. & Tuana, B. S. Cardiac-Specific Cre Induces Age-Dependent Dilated Cardiomyopathy (DCM) in Mice. Molecules 24, 1189 (2019).

29. Rashbrook, V. S., Brash, J. T. & Ruhrberg, C. Cre toxicity in mouse models of cardiovascular physiology and disease. Nat Cardiovasc Res 1, 806–816 (2022).

30. Bertero, E. & Maack, C. Calcium Signaling and Reactive Oxygen Species in Mitochondria. Circ Res 122, 1460–1478 (2018).

31. Aon, M. A. & Cortassa, S. Coherent and robust modulation of a metabolic network by cytoskeletal organization and dynamics. Biophys Chem 97, 213–231 (2002).

32. Saks, V. et al. Cardiac system bioenergetics: metabolic basis of the Frank-Starling law. J Physiol 571, 253–273 (2006).

33. Hofhuis, J. et al. Dysferlin links excitation-contraction coupling to structure and maintenance of the cardiac transverse-axial tubule system. Europace 22, 1119–1131 (2020).

34. Jung, P. et al. Increased cytosolic calcium buffering contributes to a cellular arrhythmogenic substrate in iPSC-cardiomyocytes from patients with dilated cardiomyopathy. Basic Res Cardiol 117, 5 (2022).

35. deAlmeida, A. C., van Oort, R. J. & Wehrens, X. H. T. Transverse aortic constriction in mice. J Vis Exp 1729 (2010) doi:10.3791/1729.

36. Cheillan, D. Zellweger Syndrome Disorders: From Severe Neonatal Disease to Atypical Adult Presentation. Adv Exp Med Biol 1299, 71–80 (2020).

37. Ansermet, C., et al. Renal tubular peroxisomes are dispensable for normal kidney function. JCI Insight 7, e155836 (2022).

38. Nuebel, E. et al. The biochemical basis of mitochondrial dysfunction in Zellweger Spectrum Disorder. EMBO Rep 22, e51991 (2021).

39. Thoms, S., Grønborg, S. & Gärtner, J. Organelle interplay in peroxisomal disorders. Trends in Molecular Medicine 15, 293–302 (2009).

40. Tanaka, H. et al. Peroxisomes control mitochondrial dynamics and the mitochondrion-dependent apoptosis pathway. J Cell Sci 132, jcs224766 (2019).

41. Launay, N. et al. Imbalanced mitochondrial dynamics contributes to the pathogenesis of X-linked adrenoleukodystrophy. Brain 147, 2069–2084 (2024).

42. Zemniaçak, Â. B. et al. Disruption of mitochondrial bioenergetics and calcium homeostasis by phytanic acid in the heart: Potential relevance for the cardiomyopathy in Refsum disease. Biochim Biophys Acta Bioenerg 1864, 148961 (2023).

43. Zhang, X. et al. Fasting induces hepatic lipid accumulation by stimulating peroxisomal dicarboxylic acid oxidation. Journal of Biological Chemistry 296, (2021).

44. Hachmann, M. et al. Tafazzin deficiency causes substantial remodeling in the lipidome of a mouse model of Barth Syndrome cardiomyopathy. Front Mol Med 4, 1389456 (2024).

45. Pfeiffer, K. et al. Cardiolipin stabilizes respiratory chain supercomplexes. J Biol Chem 278, 52873–52880 (2003).

46. Hu, D. et al. TMEM135 links peroxisomes to the regulation of brown fat mitochondrial fission and energy homeostasis. Nat Commun 14, 6099 (2023).

47. Tian, R. et al. Thermodynamic limitation for Ca2+ handling contributes to decreased contractile reserve in rat hearts. Am J Physiol 275, H2064–2071 (1998).

48. Tian, R., Nascimben, L., Ingwall, J. S. & Lorell, B. H. Failure to maintain a low ADP concentration impairs diastolic function in hypertrophied rat hearts. Circulation 96, 1313–1319 (1997).

49. Baumeister, P. & Quinn, T. A. Altered Calcium Handling and Ventricular Arrhythmias in Acute Ischemia. Clin Med Insights Cardiol 10, 61–69 (2016).

50. Sargsyan, Y. et al. Peroxisomes contribute to intracellular calcium dynamics in cardiomyocytes and non-excitable cells. Life Sci Alliance 4, e202000987 (2021).

51. Todt, H. et al. Oral batyl alcohol supplementation rescues decreased cardiac conduction in ether phospholipid-deficient mice. J Inherit Metab Dis 43, 1046–1055 (2020).

52. Huffnagel, I. C. et al. Rhizomelic chondrodysplasia punctata and cardiac pathology. J Med Genet 50, 419–424 (2013).

53. Dorninger, F. et al. Ether lipid transfer across the blood-brain and placental barriers does not improve by inactivation of the most abundant ABC transporters. Brain Res Bull 189, 69–79 (2022).

54. Fallatah, W. et al. A Pex7 Deficient Mouse Series Correlates Biochemical and Neurobehavioral Markers to Genotype Severity-Implications for the Disease Spectrum of Rhizomelic Chondrodysplasia Punctata Type 1. Front Cell Dev Biol 10, 886316 (2022).

55. Braverman, N. E. & Moser, A. B. Functions of plasmalogen lipids in health and disease. Biochimica et Biophysica Acta (BBA) - Molecular Basis of Disease 1822, 1442–1452 (2012).

56. Chen, T. et al. Cardioprotection from oxidative stress in the newborn heart by activation of PPARγ is mediated by catalase. Free Radical Biology and Medicine 53, 208–215 (2012).

57. Dorninger, F. et al. Overlapping and Distinct Features of Cardiac Pathology in Inherited Human and Murine Ether Lipid Deficiency. International Journal of Molecular Sciences 24, 1884 (2023).

58. Zou, Y. et al. Plasticity of ether lipids promotes ferroptosis susceptibility and evasion. Nature 585, 603–608 (2020).

59. Aldrovandi, M. & Conrad, M. Ferroptosis: the Good, the Bad and the Ugly. Cell Res 30, 1061–1062 (2020).

60. Shirakabe, A. et al. Drp1-Dependent Mitochondrial Autophagy Plays a Protective Role Against Pressure Overload–Induced Mitochondrial Dysfunction and Heart Failure. Circulation 133, 1249–1263 (2016).

61. Wang, M. et al. PEX5 prevents cardiomyocyte hypertrophy via suppressing the redox-sensitive signaling pathways MAPKs and STAT3. European Journal of Pharmacology 906, 174283 (2021).

62. Wang, M. et al. SIRT3 improved peroxisomes-mitochondria interplay and prevented cardiac hypertrophy via preserving PEX5 expression. Redox Biol 62, 102652 (2023).

63. Frey, N., Katus, H. A., Olson, E. N. & Hill, J. A. Hypertrophy of the Heart. Circulation 109, 1580–1589 (2004).

64. Toischer, K. et al. Differential Cardiac Remodeling in Preload Versus Afterload. Circulation 122, 993–1003 (2010).

65. Chaudhry, A., Shi, R. & Luciani, D. S. A pipeline for multidimensional confocal analysis of mitochondrial morphology, function, and dynamics in pancreatic β-cells. American Journal of Physiology-Endocrinology and Metabolism 318, E87–E101 (2020).

66. Berulava, T. et al. Changes in m6A RNA methylation contribute to heart failure progression by modulating translation. European Journal of Heart Failure 22, 54–66 (2020).

67. Yifrach, E. et al. Defining the Mammalian Peroxisomal Proteome. Subcell Biochem 89, 47–66 (2018).

68. Kohlhaas, M. et al. Elevated Cytosolic Na+ Increases Mitochondrial Formation of Reactive Oxygen Species in Failing Cardiac Myocytes. Circulation 121, 1606–1613 (2010).

69. Kohlhaas, M. & Maack, C. Adverse Bioenergetic Consequences of Na+-Ca2+ Exchanger– Mediated Ca2+ Influx in Cardiac Myocytes. Circulation 122, 2273–2280 (2010).

70. Helmes, M. et al. Mimicking the cardiac cycle in intact cardiomyocytes using diastolic and systolic force clamps; measuring power output. Cardiovasc Res 111, 66–73 (2016).

71. Bruegmann, T. et al. Optogenetic defibrillation terminates ventricular arrhythmia in mouse hearts and human simulations. J Clin Invest 126, 3894–3904 (2016).

72. Rössler, U. et al. Efficient generation of osteoclasts from human induced pluripotent stem cells and functional investigations of lethal CLCN7-related osteopetrosis. J Bone Miner Res 36, 1621–1635 (2021).

73. Werner, N., Nickenig, G. & Sinning, J.-M. Complex PCI procedures: challenges for the interventional cardiologist. Clin Res Cardiol 107, 64–73 (2018).

74. Hanses, U. et al. Intronic CRISPR Repair in a Preclinical Model of Noonan Syndrome-Associated Cardiomyopathy. Circulation 142, 1059–1076 (2020).

75. Kleinsorge, M. & Cyganek, L. Subtype-Directed Differentiation of Human iPSCs into Atrial and Ventricular Cardiomyocytes. STAR Protoc 1, 100026 (2020).

76. Tiburcy, M. et al. Defined Engineered Human Myocardium With Advanced Maturation for Applications in Heart Failure Modeling and Repair. Circulation 135, 1832–1847 (2017).

77. Seibertz, F., Reynolds, M. & Voigt, N. Single-Cell Optical Action Potential Measurement in Human Induced Pluripotent Stem Cell-Derived Cardiomyocytes. J Vis Exp 10.3791/61890 (2020) doi:10.3791/61890.

