## Supplemental Information for "Peroxisome protein import deficiency causes heart failure in mouse and human"

The Supplemental Information contains

- Supplemental Figures 1 to 9
- Supplemental Tables 1 and 2
- Supplemental Videos 1 to 4

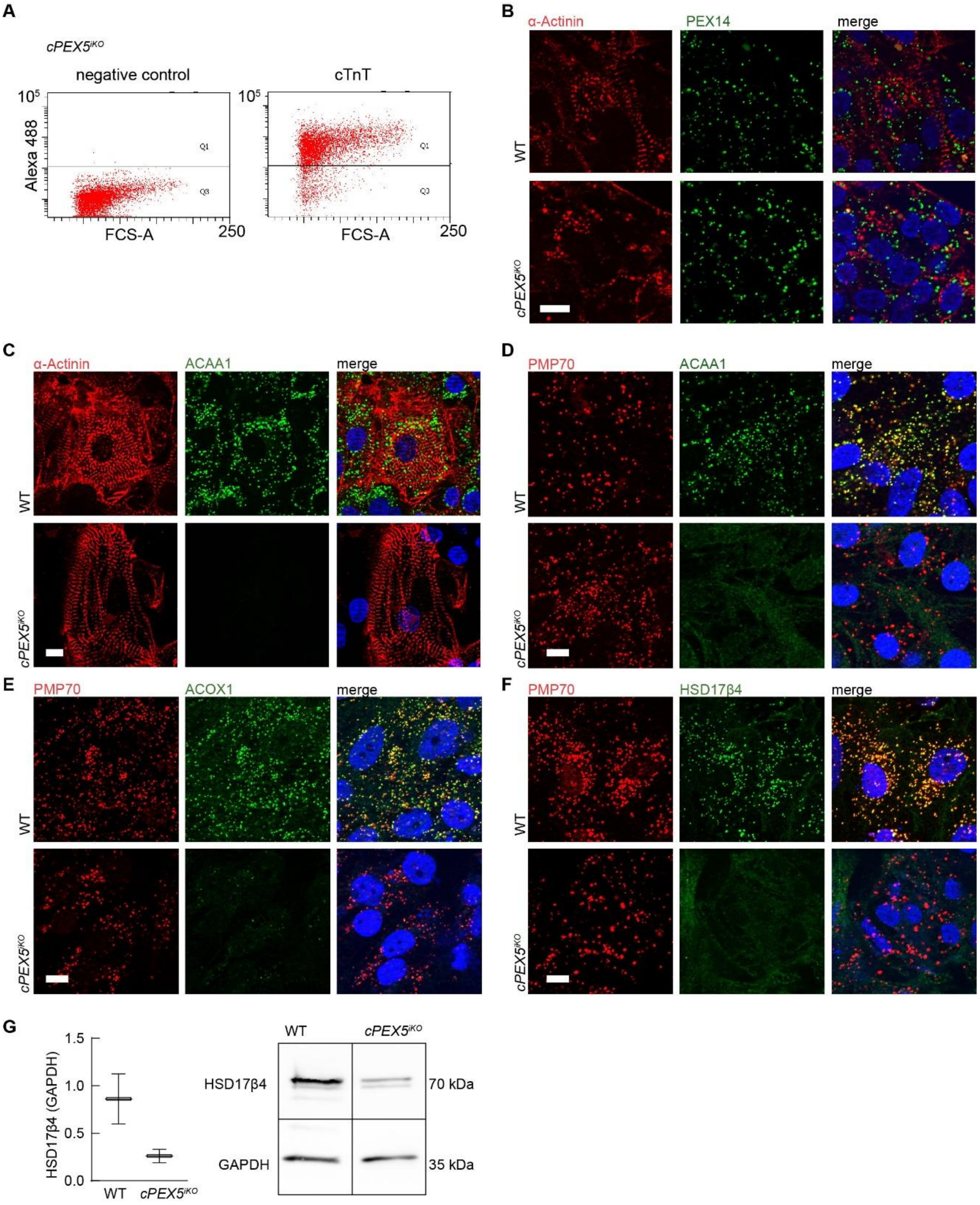


**Supplemental Figure 1: hiPSC and hiPSC-CM characterisation, rel. to Figure 1.**

(**A**) Flow cytometric analysis of cTNT stained hiPSC-CM was performed at day 60 post-cardiac differentiation. Original measurement from *cPEX5$^{iKO}$*. (**B**) Staining of peroxisomal matrix protein PEX14 shows increased size and decreased number of peroxisomes in *cPEX5$^{iKO}$*. Co-staining with α-Actinin. (**C-G**) Peroxisomal import defect was shown by co-staining of PMP70, a peroxisomal membrane protein, or sarcomeric α-Actinin with the peroxisomal matrix proteins ACAA1 (C,D), ACOX1 (E) and HSD17B4 (F) and Western blotting of the matrix protein HSD17B4 (G).

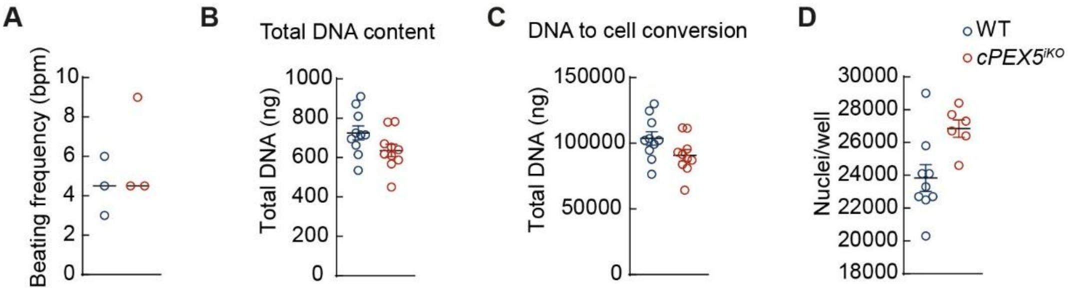


**Supplemental Figure 2: Defects mitochondrial structure in PEX5$^{iKO}$ hiPSC-CM , rel. to Figure 2**

No differences in beating frequency (**A**), total DNA content (**B**), DNA to cell conversion (**C**) or cells per well (**D**) between WT and PEX5$^{iKO}$ hiPSC-CM used for measurement of mitochondrial respiratory function.

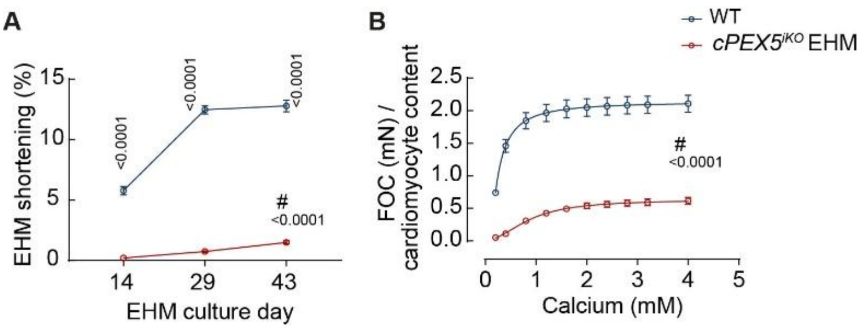


**Supplemental Figure 3: EHM characterization, rel. to Figure 3.**

EHM were generated from human WT hiPSC-CM and *cPEX5$^{iKO}$*. (**A**) Optical analysis of EHM function in 48 well plate shows reduced shortening (% change of baseline pole distance) during EHM development. (**B**) After normalization to cardiomyocyte number, force of contraction remains lower in *cPEX5$^{iKO}$*. N = 8 EHM per group. #two-way ANOVA; Bonferroni's multiple comparisons.

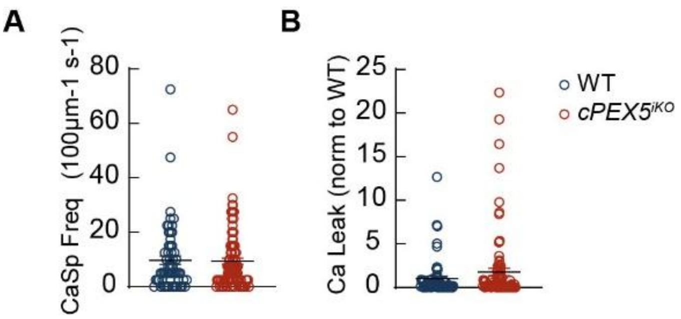


**Supplemental Figure 4: Defects in *cPEX5$^{iKO}$* calcium handling, rel. to Figure 4.**

(**A,B**) $Ca^{2+}$ spark frequency (A) and $Ca^{2+}$ leak (B) were not changed in *cPEX5$^{iKO}$* compared to WT.

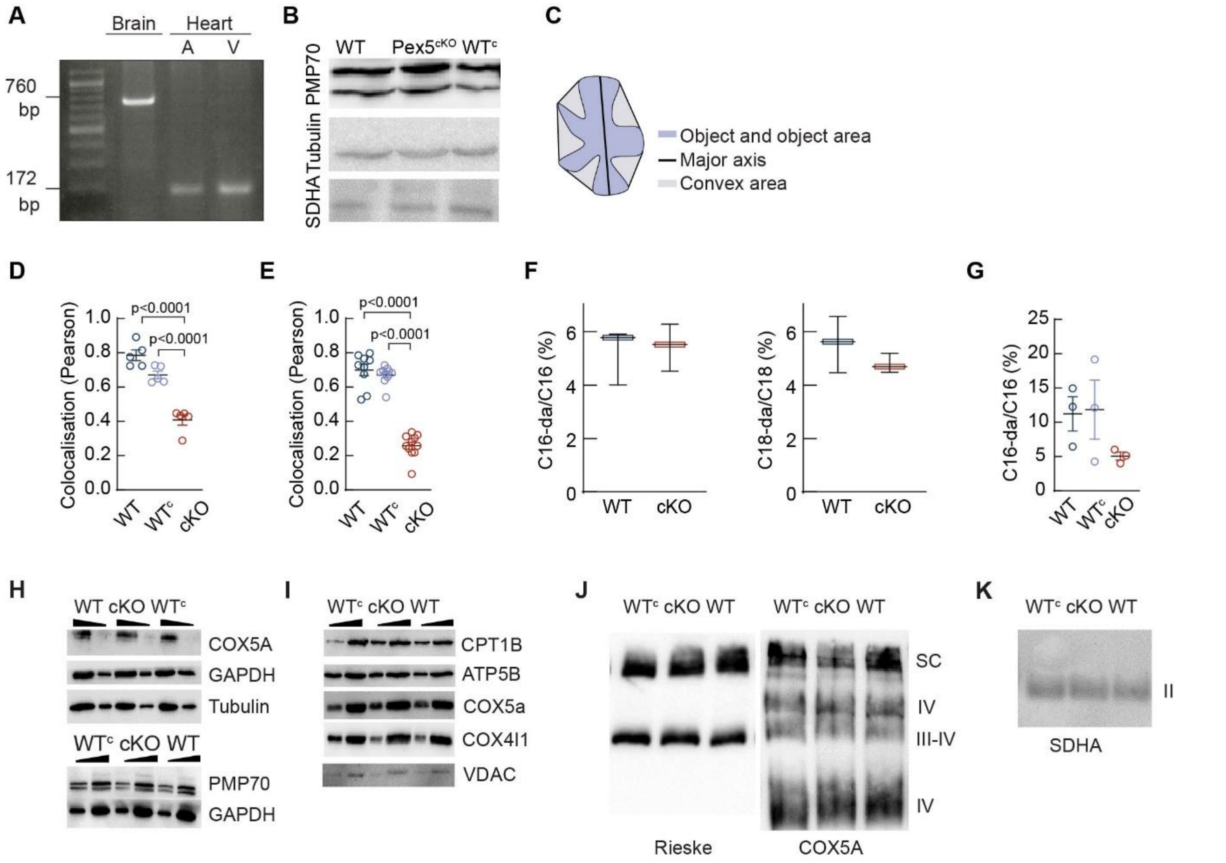


**Supplemental Figure 5: Characterization of $Pex5^{cKO}$, rel. to Fig. 5.**

(**A**) Tissue-specific knockout of *Pex5* in atria (A) and ventricles (V), but not in brain of $Pex5^{cKO}$ mice confirmed by PCR. (**B**) Equal PMP70 expression in $Pex5^{cKO}$ and WT ventricles. (**C**) Schematic of an object analysis from confocal and STED images in Figure 5F,G. Object area, major axis and convex area are used in the calculation of shape parameters of peroxisomes. (**D,E**) Loss of DAP-AT (D) and PHYH (E) in peroxisomes of isolated cardiomyocytes of fasted $Pex5^{cKO}$ mice. Quantification of Figure 5 K and M, Pearson coefficient. One-way ANOVA, Tukey's multiple comparisons. N = 5-9 cells per group. (**F**) No difference in C16 and C18 plasmalogen levels analyzed in EDTA blood samples from WT and $Pex5^{cKO}$ mice. (**G**) Non-significant reduction of C16 plasmalogens in $Pex5^{cKO}$ hearts. (**H**) Western blot analyses of cardiac tissue lysates using antibodies against mitochondrial protein COX5A and peroxisomal PMP70. Cytosolic proteins Tubulin and GAPDH were used as a control. (**I**) Isolated mitochondria were analyzed by Western blotting using indicated mitochondrial proteins CPT1B, ATP5a, COX5a and COX4I1. Vdac was used as a loading control. (**J**) Isolated mitochondria were subjected to blue native analysis to separate native respiratory chain complexes. Indicated respiratory chain complexes were analyzed by Western blot analysis against the complex III component Rieske and complex IV component COX5A. (**K**) Blue Native analysis of succinate dehydrogenase (complex II).

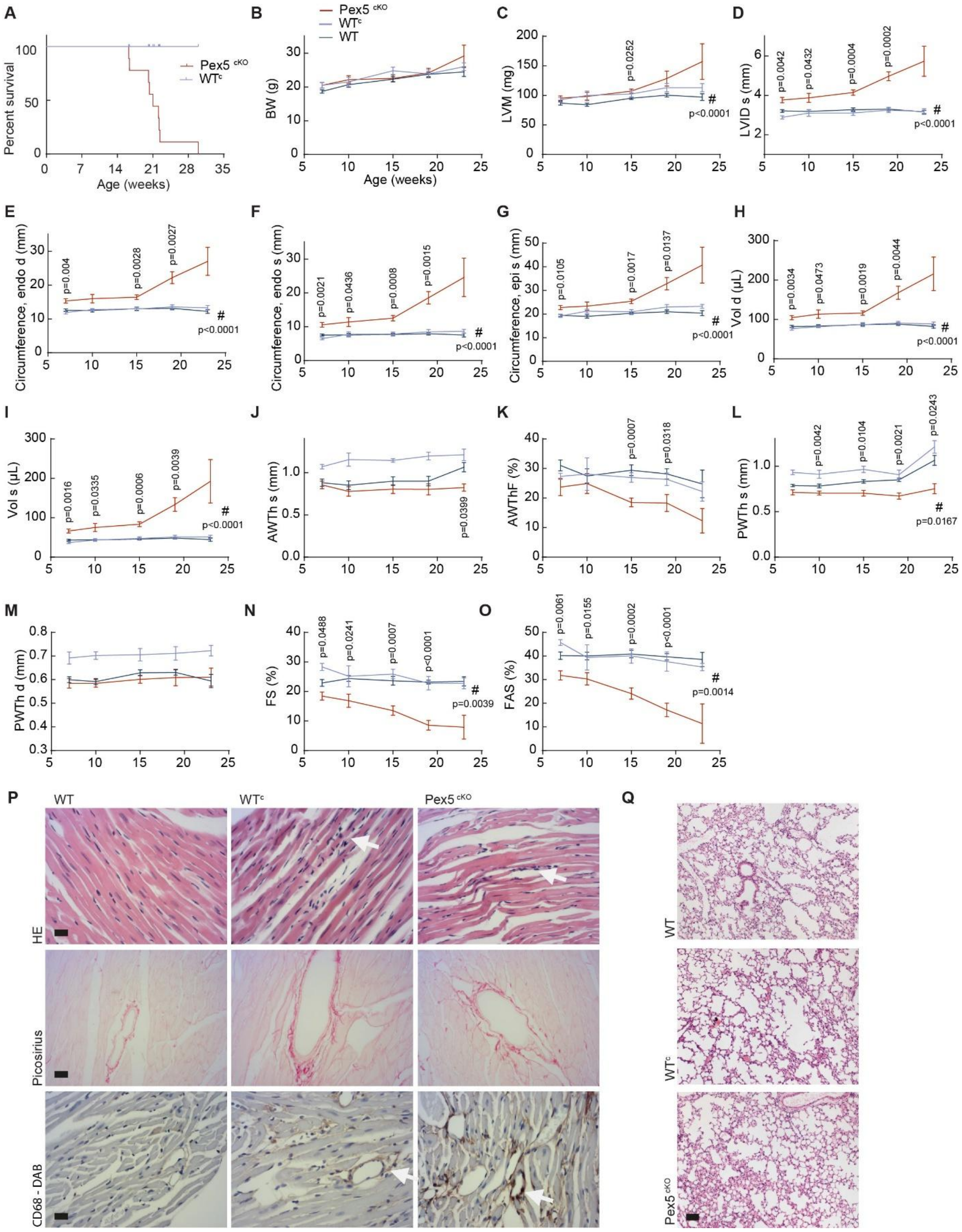

A
Percent survival
Age (weeks)
Pex5 cKO
WTc
B
BW (g)
Age (weeks)
Pex5 cKO
WTc
WT
C
LVM (mg)
p=0.0252
p<0.0001
D
LVID s (mm)
p=0.0042
p=0.0432
p=0.0004
p=0.0002
p<0.0001
E
Circumference, endo d (mm)
p=0.004
p=0.0028
p=0.0027
p<0.0001
F
Circumference, endo s (mm)
p=0.0021
p=0.0436
p=0.0008
p=0.0015
p<0.0001
G
Circumference, epi s (mm)
p=0.0105
p=0.0017
p=0.0137
p<0.0001
H
Vol d (µL)
p=0.0034
p=0.0473
p=0.0019
p=0.0044
p<0.0001
I
Vol s (µL)
p=0.0016
p=0.0335
p=0.0006
p=0.0039
p<0.0001
J
AWTh s (mm)
p=0.0399
K
AWThF (%)
p=0.0007
p=0.0318
L
PWTh s (mm)
p=0.0042
p=0.0104
p=0.0021
p=0.0243
p=0.0167
M
PWTh d (mm)
N
FS (%)
p=0.0488
p=0.0241
p=0.0007
p<0.0001
p=0.0039
O
FAS (%)
p=0.0061
p=0.0155
p=0.0002
p<0.0001
p=0.0014
P
WT
WTc
Pex5 cKO
HE
Picosirius
CD68 - DAB
Q
WT
WTc
Pex5 cKO

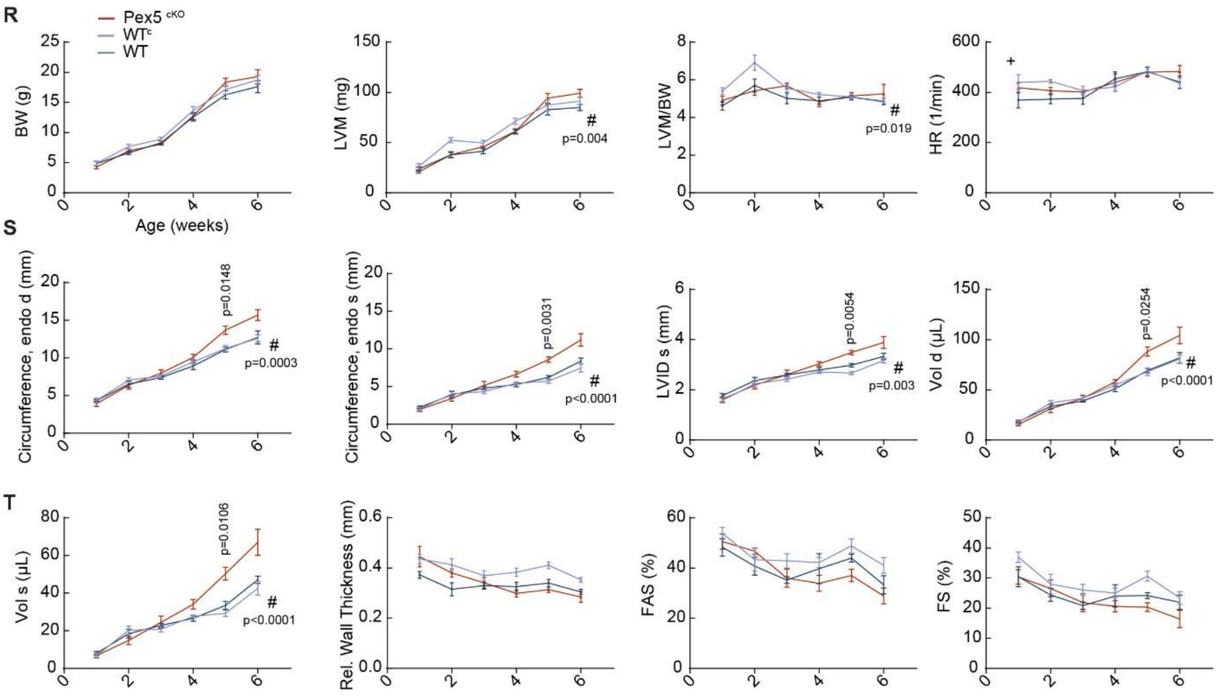


**Supplemental Figure 6: Characterization of Pex5$^{cKO}$ heart function, rel. to Fig. 6.**

(**A**) Kaplan Maier survival curve shows early death of Pex5$^{cKO}$ mice, but Cre-expressing WT$^{c}$ mice show normal life span. p<0.01. N = 5-7 mice per group. (**B-O**) Echocardiography of 7- to 23-week old mice shows dilated hearts with decreased contraction indicating severe DCM in Pex5$^{cKO}$ mice. N = 3-16 mice per group and time point, one-way ANOVA, Tukey's multiple comparisons. (**P**) Picrosirius staining shows no signs of fibrosis in Pex5$^{cKO}$ hearts. HE- and CD68-DAB staining (highly expressed by macrophages) of heart tissue sections show inflammation and macrophage infiltration (white arrows) in Pex5$^{cKO}$ and WT$^{c}$ mouse hearts. (**Q**) No signs of pathology in HE-staining of Pex5$^{cKO}$ and WT$^{c}$ lung tissue. Scales: 20 µm. (**R-T**) Echocardiography of 1- to 6-week old mice shows no functional differences in the first 6 weeks of age, but first signs of dilation in 4 week old Pex5$^{cKO}$ mice. N = 5-7 mice per group and time point, one-way ANOVA, Tukey's multiple comparisons BW = body weight, LVW = left ventricular weight, HR = heart rate, AWThd = anterior wall thickness diastole, AWThs = anterior wall thickness systole, PWThd = posterior wall thickness diastole, PWThs = posterior wall thickness systole, LVIDd = left ventricular internal diameter end diastole, LVIDs = left ventricular internal diameter end systole, FS = fractional shortening, FAS = fractional area shortening, AWThF = anterior wall thickening fraction, PWThF = posterior wall thickening fraction, Vol s = volume systole, Vol d = volume diastole, Epi = epicardium.

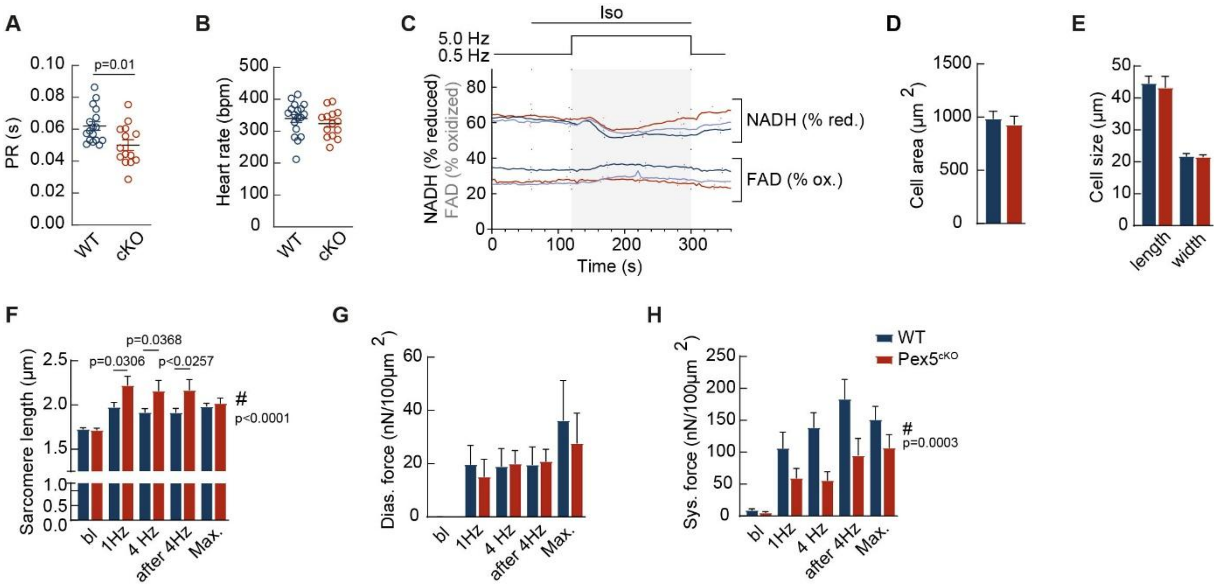


**Supplemental Figure 7: Characterization of $Pex5^{cKO}$ cardiac mitochondrial energentics and force production, rel. to Fig. 7.**

(**A,B**) Statistical analysis of the PR interval, 50.1 ± 0.3 ms vs. 63.7 ± 0.3 ms (A) and the spontaneous beating rate (B) in cardiac electrograms of explanted hearts from WT and $Pex5^{cKO}$. (**C**) Autofluorescences of NAD(P)H and FAD reveal a trend towards a more reduced redox status of $Pex5^{cKO}$ compared to WT cells. (**D-H**) Force measurement in unloaded cardiomyocytes. Similar cell area (D) and size (E) during strech were aquired to ensure that the cells were streched equally and force is therefore theoretically aquired from the same amounts of sarcomers. Cell length corresponds to the distance between the to holders that strech the cell and measure the force. $Pex5^{cKO}$ cells show increased sarcomere length (F). Diastolic force (G) is not changed, whereas systolic force (H) is significantly decreased in $Pex5^{cKO}$ myocytes. Two-way ANOVA, Bonferroni's multiple comparisons.

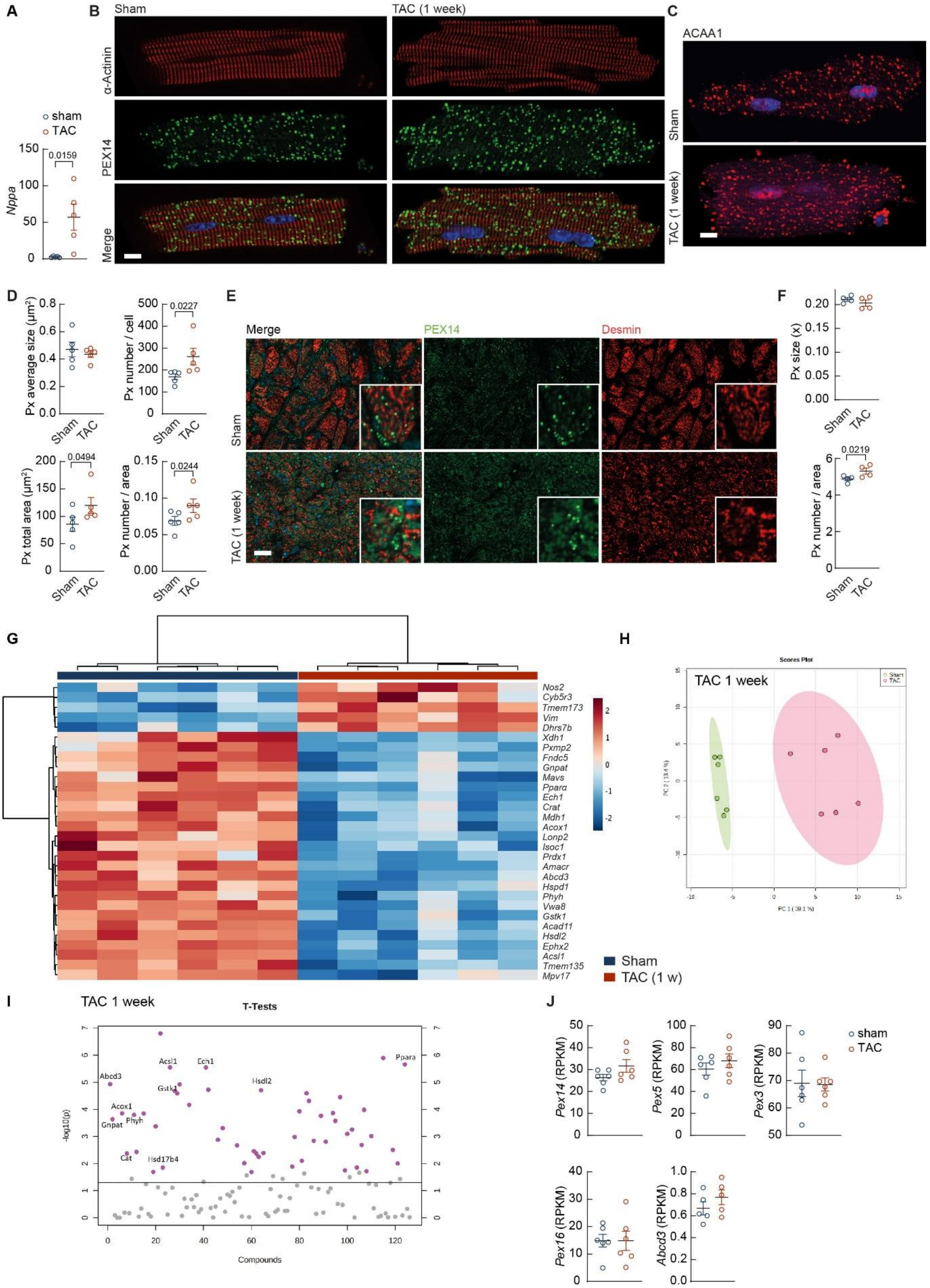

A
B
Sham
TAC (1 week)
C
ACAA1
α-Actinin
PEX14
Merge
sham
TAC
Nppa
0.0159
Sham
TAC (1 week)
D
Px average size (μm²)
Px number / cell
0.0227
Px total area (μm²)
0.0494
Px number / area
0.0244
E
Merge
PEX14
Desmin
F
Px size (x)
Px number / area
0.0219
G
Nos2
Cyb5r3
Tmem173
Vim
Dhrs7b
Xdh1
Pxmp2
Fndc5
Gnpat
Mavs
Pparα
Ech1
Crat
Mdh1
Acox1
Lonp2
Isoc1
Prdx1
Amacr
Abcd3
Hspd1
Phyh
Vwa8
Gstk1
Acad11
Hsdl2
Ephx2
Acsl1
Tmem135
Mpv17
Sham
TAC (1 w)
H
Scores Plot
TAC 1 week
I
TAC 1 week
T-Tests
Abcd3
Acsl1
Ech1
Ppara
Gstk1
Hsdl2
Acox1
Gnpat
Phyh
Cat
Hsd17b4
-log10(p)
Compounds
J
Pex14 (RPKM)
Pex5 (RPKM)
Pex3 (RPKM)
sham
TAC
Pex16 (RPKM)
Abcd3 (RPKM)

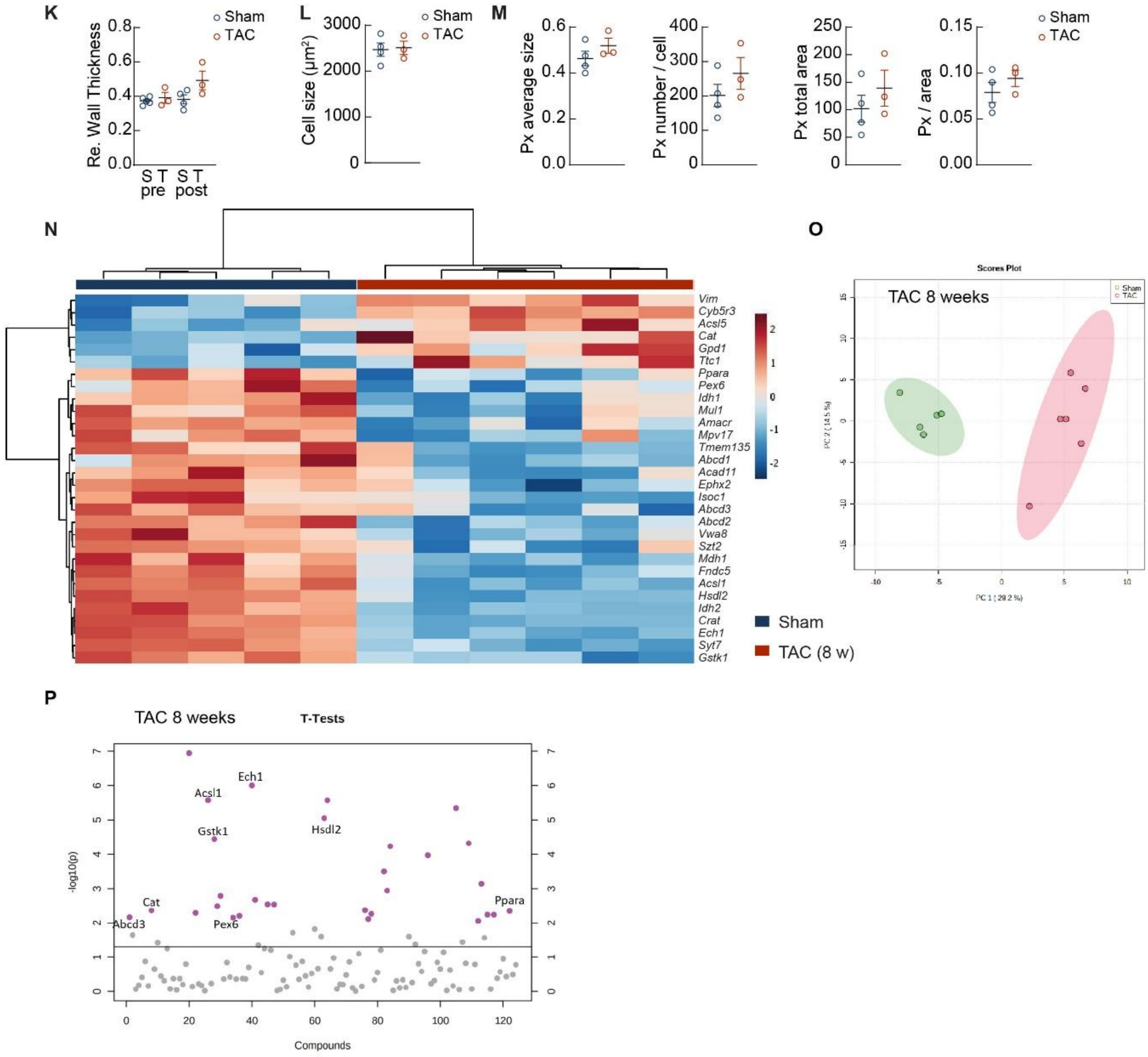


**Supplemental Figure 8: Peroxisome abundance and compositipon in the failing heart, rel. to Fig. 8.**

(**A**) *Nppa* mRNA expression significantly increased one week after TAC. qPCR. (**B,C**) Peroxisome distribution in isolated CM from sham-operated mouse hearts and mouse hearts isolated 1 week after transverse aortic constriction (TAC). Staining of PEX14 and α-Actinin (B) or ACAA1 (C) in isolated CM 1 week after TAC. (**D**) Analysis of ACAA1-stained peroxisomes (Px) from (C). N = 4-5 mice per group. (**E**) Paraffin-embedded sections stained with anti-PEX14 and anti-Desmin. Scale: 10 µm. (**F**) Peroxisome number per area increased significantly 1 week after TAC, peroxisome size did not change. Quantification of (E). N = 4 mice. (**G-I**) RNA sequencing one week after TAC. Heatmap (G), PCA plot (H) and (I) T-tests. (**J**) *Pex14, Pex5, Pex3*, *Pex16* and *Abcd3* mRNA expression unchanged one week after TAC. Data from RNA sequencing. (**K**) Echocardiographic analysis of WT mice before (pre) and 8 weeks after TAC (post) shows no significant change in relative wall thickness; N=4 WT and 3 *Pex5*$^{cKO}$ mice. S = sham, T = TAC. (**L,M**) Analysis of peroxisome staining of isolated CM from sham and 8 week-TAC mouse hearts. Peroxisomes were stained by anti-PEX14 (L) or anti-ACAA1 (M). No change in cell size, peroxisome size or peroxisome number was observed. N = 4 WT, 3 TAC mice. (**N-P**) RNA sequencing 8 weeks after TAC. (N) Heatmap, (O) PCA plots and (P) T-tests.

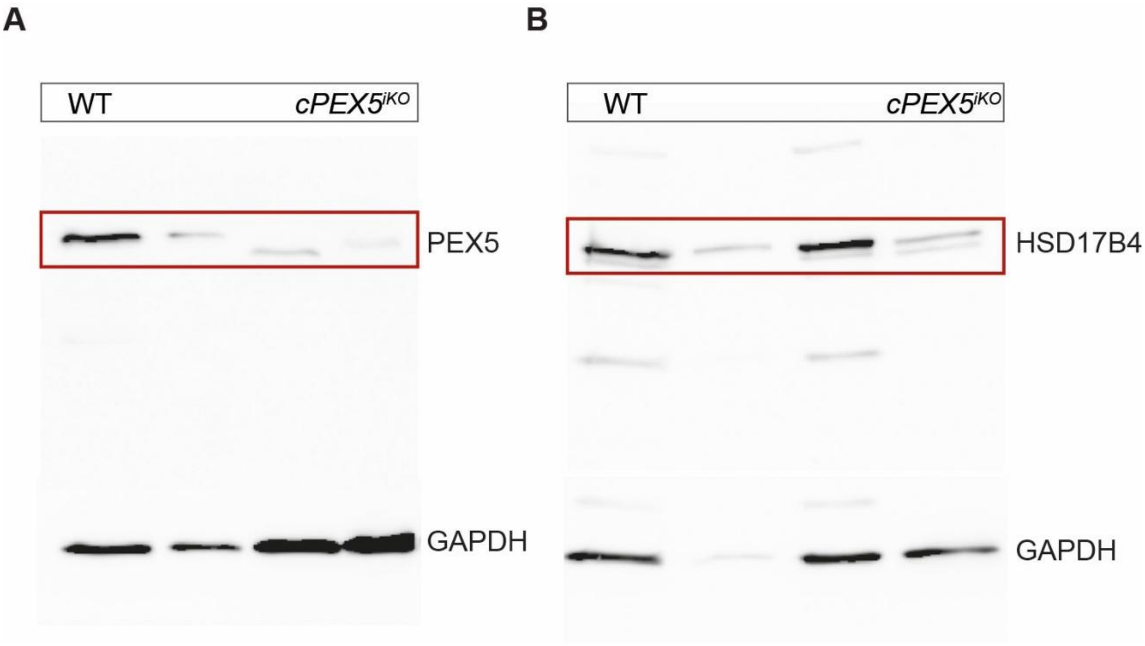


**Supplemental Figure 9: Uncropped Western Blots.**

(**A**) PEX5 expression, Fig.1 L (**B**) HSD17β4, Fig. S1G.

**Supplemental Table 1: Antibodies.**

| First antibody | Firma | Order No. | Dilution |
|---|---|---|---|
| PEX14 | Proteintech | 10594-1-AP | 1:500, 1:50 (IHC) |
| ACAA1 | Proteintech | 12319-2-AP | 1:500 |
| PMP70 | Sigma | SAB4200181 | 1:250 |
| Desmin | Progen | 10519 | 1:100 (IHC) |
| CD68 | Dako | 110876 | 1:200 |
| CAV3 | BD | 610420 | 1:250 |
| RYR2 | Thermo | MA3-916 | 1:200 |
| PHYH | Santa Cruz | sc-376727 | 1:100 |
| DAP-AT | Proteintech | 14931-1-AP | 1:100 |
| HSD17B4 | Proteintech | 15116-1-AP | 1:100, 1:100 (WB) |
| ACOX1 | Proteintech | 10957-1-AP | 1:100 |
| α-Actinin | Sigma | A7811 | 1:500 |
| CAV3 | BD | 610420 | 1:250 |
| RYR2 | Thermo | MA3-916 | 1:200 |
| OCT | BD Biosciences | 560329 | 1:100 |
| NANOG | Thermo | MA1-017 | 1:200 |
| TRA1-60 | BD Biosciences | 560173 | 1:100 |
| PEX5 | Novus Biologicals | NBP2-38443 | 1:500 |
| GAPDH | Millipore | MAB374 | 1:10000 |
| Anti-cTnT | Invitrogen | MA5-12960 | 1:500 |
| Anti-mouse IgG-Alexa Fluor® 633 | Invitrogen | A-21053 | 1:1000 |
| Anti-rabbit IgG-Alexa Fluor® 488 | Life technologies | A-11008 | 1:1000 |
| Anti-mouse IgG-Alexa Fluor® 488 | Invitrogen | A-21202 | 1:1000 |
| Anti-mouse IgG-CyTM3 | Jackson ImmunoResearch | 715-166-150 | 1:1000 |
| Star 635P rabbit | Abberior | ST635P | 1:1000 |
| Star 580 mouse | Abberior | ST580 | 1:1000 |
| ECL Anti-Rabbit IgG, HRP | Jackson ImmunoResearch | 111-035-003 | 1:10000 |
| ECL Anti-Mouse IgG, HRP | Jackson ImmunoResearch | 715-035-150 | 1:10000 |

**Supplemental Table 2: Primers used for qPCR** (Murine genes).

| Primers | Sequence for | Sequence rev | Product size (bp) |
|---|---|---|---|
| *Anp* | TCGTCTTGGCCTTTTGGCT | TCCAGGTGGTCTAGCAGGTTCT | 106 |
| *Pparα* | TCGGCGAACTATTCGGCTG | GCACTTGTGAAAACGGCAGT | 106 |
| *Cat* | CCGACCAGGGCATCAAAA | GAGGCCATAATCCGGATCTTC | 74 |
| *Phyh* | TGCCAAGTGATAGGATGGTTTC | TCAGAATCTCGGGGAGAAGG | 87 |
| *Gnpat* | GGAGTCCACCACTGGATTGT | GCAGTGATACAGGGTCTCCG | 92 |
| *Pex11a* | ACTGGCCGTAAATGGTTCAGA | CGGTTGAGGTTGGCTAATGTC | 119 |
| *Hsdl2* | CTGTTCCACTGCCACCAAGT | ATCCCGCTAGCTTCCCAGT | 70 |
| *Acsl1* | TCCTACAAAGAGGTGGCAGAACT | GGCTTGAACCCCTTCTGGAT | 68 |
| *Pln* | TGCCCAGCTAAGCTCCCATA | TTTGTTGTGCAGACTGAAGCG | 116 |
| *Gapdh* | GAAGATGGTGATGGGATT | GAAGGTGAAGGTCGGAGT | 89 |
| *Gusb* | CACGGCGATGGACCCAAGAT | CCCATTCACCCACACAACTGC | 86 |

**Supplemental Video 1. Contraction of WT EHM.** Representative videos of spontaneously beating EHM generated from WT hiPSC-CM. Related to Figure 3.

**Supplemental Video 2. Contraction of *PEX5*$^{iKO}$ EHM.** Representative videos of spontaneously beating EHM generated from c*PEX5*$^{iKO}$. Related to Figure 3.

**Supplemental Video 3. Optical Mapping of WT EHM.**

Representative video of optical mapping of WT EHM. Colors representing the time point of maximal signal amplitude after electrical point stimulation (see Figure 4I). Stimulation electrode is situated in the inferior left corner of the tissue. Related to Figure 4.

**Supplemental Video 4. Optical Mapping of *PEX5*$^{iKO}$ EHM.**

Representative videos of optical mapping of PEX5$^{iKO}$ EHM. Colors representing the time point of maximal signal amplitude after electrical point stimulation (see Figure 4I). Stimulation electrode is situated in the inferior left corner of the tissue. Related to Figure 4.
